# Emergent memory in cell signaling: Persistent adaptive dynamics in cascades can arise from the diversity of relaxation time-scales

**DOI:** 10.1101/247684

**Authors:** Tanmay Mitra, Shakti N. Menon, Sitabhra Sinha

## Abstract

The mitogen-activated protein kinase (MAPK) signaling cascade, an evolutionarily conserved motif present in all eukaryotic cells, is involved in coordinating critical cell-fate decisions, regulating protein synthesis, and mediating learning and memory. While the steady-state behavior of the pathway stimulated by a time-invariant signal is relatively well-understood, we show using a computational model that it exhibits a rich repertoire of transient adaptive responses to changes in stimuli. When the signal is switched on, the response is characterized by long-lived modulations in frequency as well as amplitude. On withdrawing the stimulus, the activity decays over timescales much longer than that of phosphorylation-dephosphorylation processes, exhibiting reverberations characterized by repeated spiking in the activated MAPK concentration. The long-term persistence of such post-stimulus activity suggests that the cascade retains memory of the signal for a significant duration following its removal, even in the absence of any explicit feedback or cross-talk with other pathways. We find that the molecular mechanism underlying this behavior is related to the existence of distinct relaxation rates for the different cascade components. This results in the imbalance of fluxes between different layers of the cascade, with the repeated reuse of activated kinases as enzymes when they are released from sequestration in complexes leading to one or more spike events following the removal of the stimulus. The persistent adaptive response reported here, indicative of a cellular “short-term” memory, suggests that this ubiquitous signaling pathway plays an even more central role in information processing by eukaryotic cells.

## I INTRODUCTION

Intra-cellular signaling networks are paradigmatic examples of complex adaptive systems that exhibit a rich repertoire of responses to stimuli [1]. Such networks mediate the response of a cell to a wide variety of extra-and intra-cellular signals primarily through a sequence of enzyme-substrate biochemical reactions [2, 3]. While the complexity of the entire signaling system is daunting [4], it is possible to gain an insight into how it functions by focusing on a key set of frequently occurring motifs. These often take the form of linear signaling cascades, referred to as pathways. One of the best known of these pathways is the mitogen-activated protein kinase (MAPK) cascade that is present in all eukaryotic cells [5, 6]. It is involved in regulating a range of vital cellular functions, including proliferation and apoptosis [6], stress response [7] and gene expression [8]. This signaling module comprises a sequential arrangement of three protein kinases, viz., MAPK, MAPK kinase (MAP2K) and MAPK kinase kinase (MAP3K). Modular function is initiated when extracellular signals stimulate membrane-bound receptors upstream of the cascade, with the information being relayed to MAP3K by a series of intermediaries. Activated kinases in each layer of the module function as enzymes for phosphorylating (and thereby activating) the kinase in the level immediately downstream, with the subsequent deactivation being mediated by corresponding dephosphorylating enzymes known as phosphatases (P’ase). The terminal kinase in this cascade, i.e., MAPK, transmits the signal further downstream by phosphorylating various proteins including transcription regulators [9]. Extensive investigations into the steady-state behavior of the cascade have contributed towards an in-depth understanding of several emergent features including ultrasensitivity [10], and oscillations [11, 12] that arise through retrograde propagation of activity [13, 14]. One of the striking features of the cascade is the occurrence of bistability which allows the system to switch between two possible states corresponding to low and high activity [12, 15–18]. This provides a post-transcriptional mechanism for obtaining a sustained response from transient signals, i.e., cellular memory [19, 20].

Memory can be understood as long-term alterations in the state of a system in response to environmental changes, which allow the system to retain information about transient signals long after being exposed to them [19]. This can arise in the cell through mechanisms such as auto-regulatory transcriptional positive feedback [21] and nucleosomal modifications [22]. In the context of cell-fate determination, it has been shown that an irreversible biochemical response can be generated from a short-lived stimulus through feedback-based bistability [20]. This corresponds to a permanent alteration of the state of the system, thereby actively maintaining ‘memory’ of the signal. As bistability has also been observed to arise through multi-site phosphorylation in signaling modules, protein phosphorylation has been suggested as a plausible post-transcriptional mechanism for cellular memory [19, 23, 24]. In particular, there have been extensive investigations of the MAPK cascade as it integrates a large range of signals received by the cell in order to control numerous cellular decisions [25–30]. While these investigations have considered the steady state behavior of the system, one may also observe transitory modulations in the response of the cascade in a changing environment. The latter could encode information about prior stimuli to which the system was exposed, and can be a potential mechanism for imparting a form of “short-term” memory to the signaling cascade.

In this paper we show that a linear MAPK cascade can indeed exhibit short-term memory through transient modulations in its response to an environmental change. Crucially, this can arise even in the absence of explicit feedback between different layers or cross-talk with other pathways. These modulations can persist long after the initial trigger, lasting for durations that are several orders of magnitude longer than the time-scales associated with phosphorylation-dephosphorylation processes. We demonstrate that this occurs both when a signal begins activating the MAPK cascade, as well as when it is withdrawn. On application of the stimulus, the module exhibits long-lived frequency and amplitude modulations in the activation profile of the constituent kinases. Following the withdrawal of stimulus, activity in the cascade decays over an extremely long time-scale, during which reverberatory dynamics, characterized by large-amplitude spiking in MAP Kinase activity, can be observed. We explain the emergence of such long-lived memory of the withdrawn stimulus in terms of the imbalance of fluxes between different layers of the cascade, which results from the diversity of relaxation time-scales of the cascade components, and the reuse of activated kinases as enzymes when they are released from sequestration. This phenomenon is seen to be robust with respect to variations in the molecular concentrations of the constituent kinases and phosphatases. Our results reveal that a biochemical signaling module as simple as the MAPK cascade is capable of exhibiting short-term memory that is manifested as persistent modulations in the adaptive response of the system to changes in stimuli.

## II. METHODS

The dynamics of the three layer MAPK signaling cascade has been simulated using the Huang-Ferrell model [10]. Each of the constituent kinase and phosphatase-mediated enzyme-substrate reactions comprise (i) a reversible step corresponding to the formation of the enzyme-substrate complex and (ii) an irreversible product formation step corresponding to the activation/deactivation of a kinase, as described in the Supplementary Information. The time-evolution of the molecular concentrations of the different components of the cascade are modeled using a set of coupled ordinary differential equations (see Supplementary Information) that are integrated using the stiff solver ode15s implemented in *MATLAB Release 2010b*. Note that the quasi-steady-state hypothesis has not been invoked [31]. To ensure that initially all kinases are non-phosphorylated we prepare the initial resting state of the system by simulating it for a long duration (∼ 10^6^ mins) in the absence of any signal. Subsequently MAP3K is exposed to a stimulus of amplitude *S* and duration 5000 minutes. Following the removal of the stimulus, we continue to simulate the system until it returns to the resting state or the simulation duration exceeds 10^4^ minutes.

We have analyzed the long-lived reverberatory activity of the cascade after the removal of the stimulus by using the following measures:

i. *The primary recovery time* (*τ*_PR_). Following the activation of the cascade by introducing a stimulus, the maximum concentration *R*_max_ of MAPK^**^ is recorded (Note that ** represents a doubly phosphorylated kinase while * indicates that it is singly phosphorylated). On removing the stimulus, MAPK activity starts to decay. The time taken for MAPK** to monotonically decrease to half of *R*_max_ is defined as the primary recovery time (*τ*_PR_).
ii. *Number of spikes during relaxation* (*N_r_*). Following primary recovery, MAPK activity may exhibit a series of spikes, which are defined to be occurring whenever MAPK^**^ concentration exceeds 70% of *R*_max_. The num-ber of such spikes that are observed before the cascade reaches its resting state is designated as *N_r_*.
iii. *The total duration of reverberatory activity* (*τ_r_*). When spiking is observed in MAPK activity following the removal of the applied stimulus, the reverberatory activity duration is defined as the interval between the termination of primary recovery and the final spike event, i.e., *τ_r_*= *t_final_* − *τ_PR_*. The time of the *i*th spike *t_i_* is defined as the instant when MAPK activity reaches maximum during that particular event. For *τ_PR_* > 6000 mins, the total duration of the reverberatory activity may not be measured accurately as the total simulation duration does not exceed 10^4^ minutes.
iv. *The total memory time* (*τ_m_*). The total duration of memory activity following removal of the applied stimulus is defined as the sum of the primary recovery time and the total duration of reverberatory activity, i.e., *τ_m_* = *τ_PR_*+*τ_r_*. Note that when the steady-state behavior of the cascade in presence of the signal is oscillatory, on withdrawing the signal the activity may decay extremely rapidly resulting in *τ_m_* ≈ 0.
v. *Relaxation time* (*τ*_x_). For the situations where the steady state corresponds to a fixed-point attractor we define a relaxation time *τ*_x_ for each constituent of the cascade. This is the time required by its concentration to evolve to the half-way point between the resting state and steady state values.

## III RESULTS

*Emergence of persistent modulations in kinase activity.* For the results reported in this paper we consider the Huang-Ferrell model of the MAPK signaling cascade [10], schematically shown in Fig. 1(a). Typically, investigations into the dynamics of this model focus on the steady-state response to sustained stimulation. In contrast, here we report on the transient activity of the system following a change in the stimulus. Specifically, we describe the response immediately following the introduction of a signal of amplitude *S* and that following its removal. Our results reveal that such transients can be unexpectedly long-lived, lasting for durations that are much longer compared to the time-scales associated with the phosphorylation and dephosphorylation processes in the cascade (Fig. 1, b).

**FIG. 1:**
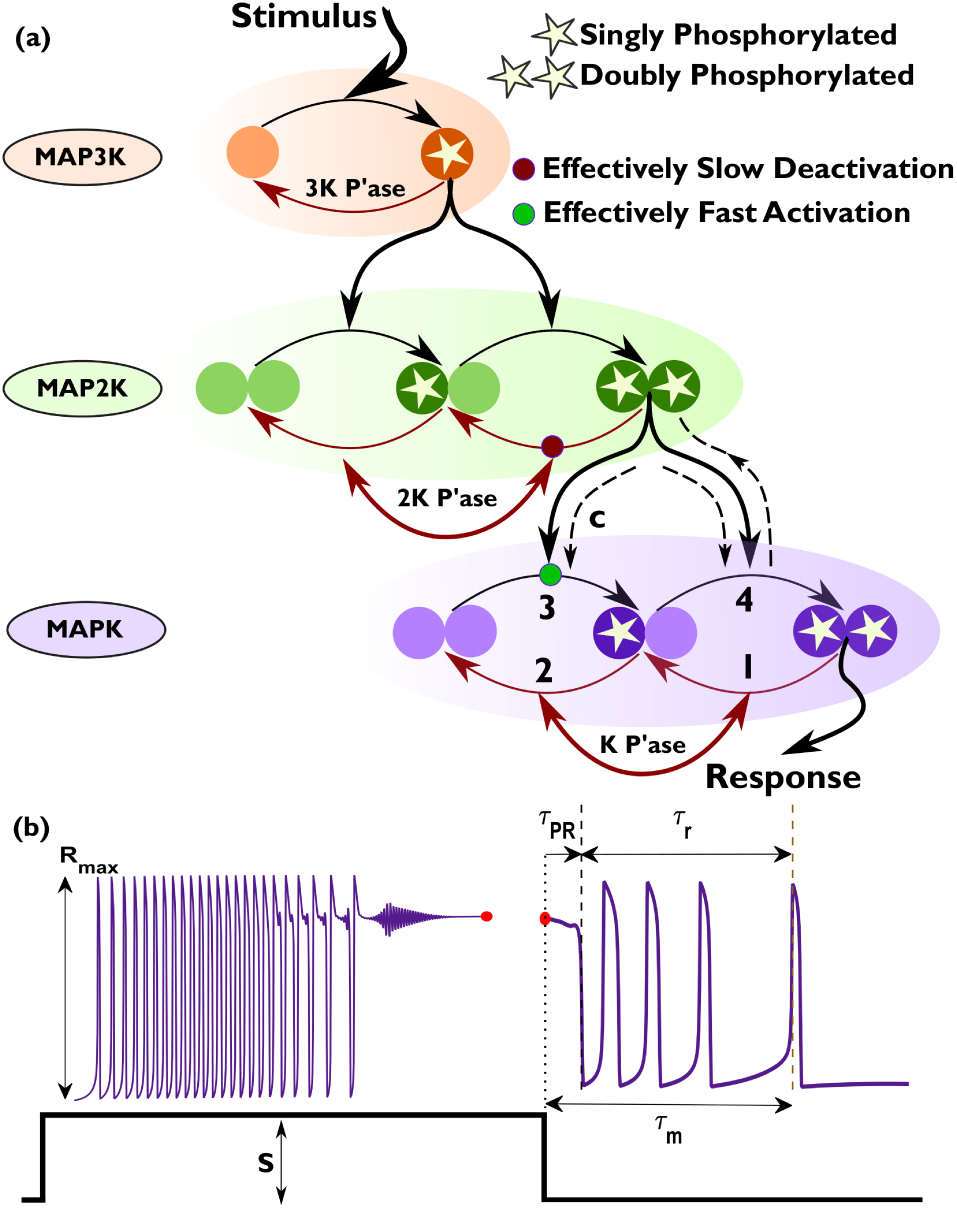
Adaptive response of MAPK cascade to a changing stimulus. (a) Schematic representation of a linear MAPK cascade comprising three layers. Signaling is initiated by a stimulus S activating MAPK kinase kinase (MAP3K). Activation/deactivation of kinases is achieved by adding/removing phosphate groups, which is referred to as phosphorylation/dephosphorylation respectively. The activated MAP3K regulates the phosphorylation of MAPK kinase (MAP2K). Doubly phosphorylated MAP2K, in its turn, controls the activation of MAPK. The response of the cascade to the signal is measured in terms of MAPK activity, viz., the concentration of doubly phosphorylated MAPK. Deactivation of a phosphorylated kinase is regulated by the corresponding phosphatase (indicated by P’ase) in the corresponding layer of the cascade. The numbers 1 − 4 represent the sequence of events that lead to the emergence of a large amplitude spiking response following the withdrawal of the stimulus. The enzyme-substrate protein complex formed during activation of MAPK by doubly phosphorylated MAP2K is indicated by “c”. Broken lines have been used to highlight the principal processes that drive the reverberatory dynamics, which functions as a memory of the signal (see text for details). (b) Schematic illustrating the emergence of long-lived transient modulations of MAPK activity in response to initiation of a signal of optimal strength *S*. Withdrawing the stimulus can result in persistent large-amplitude spiking in the response of MAPK, suggestive of a form of “short-term” memory. The maximum response of MAPK to the stimulus is denoted by *R*_max_. The primary recovery time (*τ_PR_*) is characterized as the duration following withdrawal of stimulus after which MAPK activity decreases to its half-maximum value (*R*_max_/2) for the first time. The duration over which reverberatory dynamics occurs is indicated by *τ_r_*, while the total duration for which memory of the withdrawn stimulus persists is 
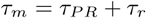

We first report the behavior of a cascade that is initially in the resting state (characterized by the absence of any phosphorylated components) when it is exposed to a signal. The transient activity that immediately follows exhibits several non-trivial features such as regular spiking in the activity of MAP2K and MAPK depending on the total concentrations of the kinases (Fig. 2, b-e) and the signal strength. For a fixed initial state and signal strength, the spikes can further show modulation in their frequency (Fig. 2, c-e) as well as amplitude (Fig. 2, b and d). In certain cases, both types of modulation can be observed (Fig. 2, d). In the representative time series of MAPK activity shown in Fig. 2(a-e), the system dynamics eventually converges to a stable fixed point (Fig. 2, a-d) or a stable limit cycle (Fig. 2, e), with the attractors being independent of initial conditions (corresponding phase space projections are shown in Fig. 2, f-i). Note that when phosphorylated components are initially present, the system reaches the asymptotic state faster.

**FIG. 2:**
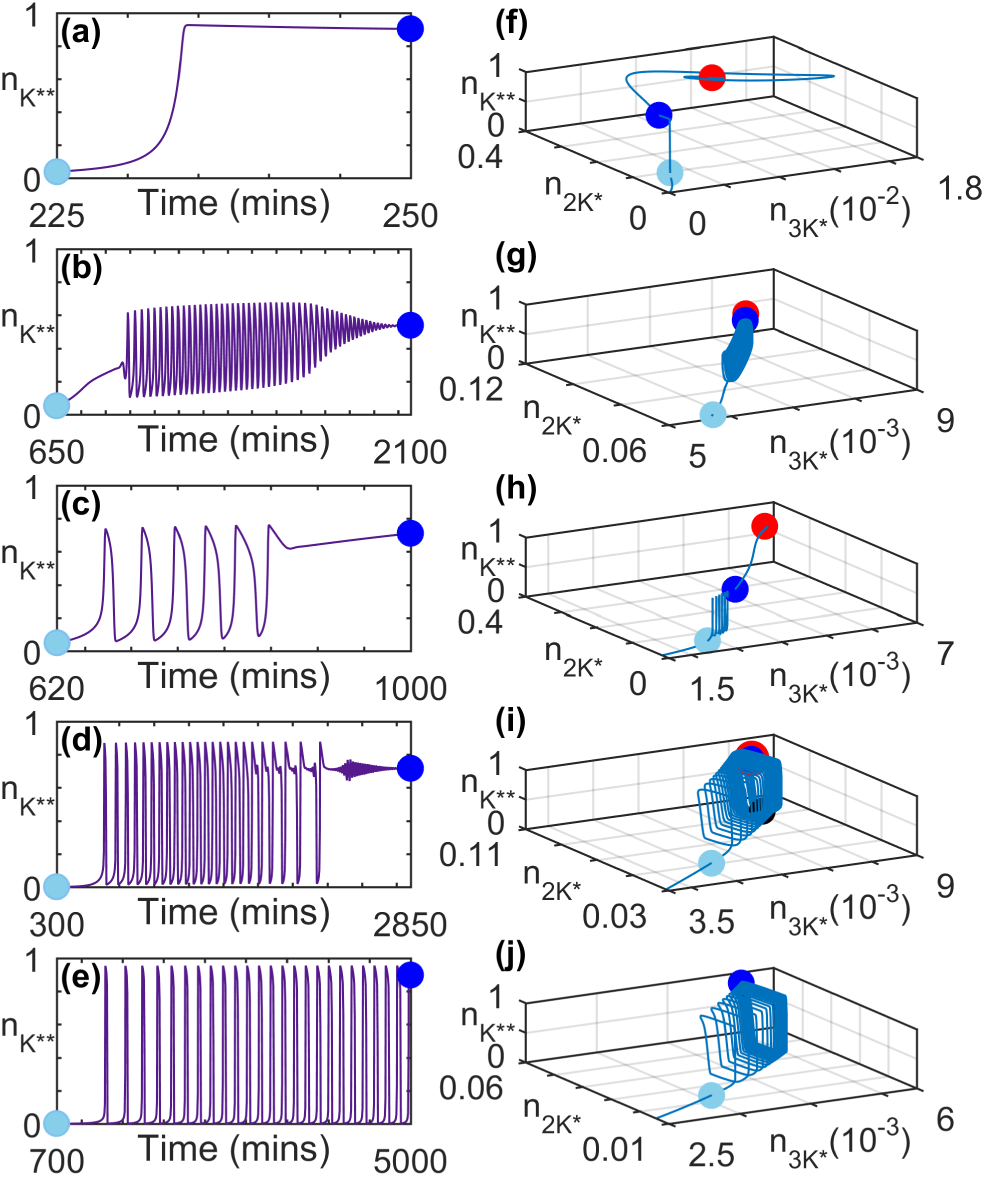
Transient activity in MAPK cascade immediately following the application of a stimulus having amplitude *S* = 1.2 × 10^−6^*µM* at *t* = 0. (a-e) Characteristic time series for the normalized concentration of doubly phosphorylated MAPK (*n_K_*\*\*) shown for different total concentrations of kinases. The concentration of active MAPK is insignificant prior to the time periods shown in panels (a-e). (f-j) Trajectories representing the evolution of the systems in panels (a-e) in the projection of the phase-space on the planes comprising normalized concentrations of active MAP3K (*n_3K_*\*), singly phosphorylated MAP2K (*n_2K_*\*) and active MAPK (*n_K_*\*\*). The concentrations have been normalized by the total concentration of MAP3K ([3*K*]_*tot*_), MAP2K ([2*K*]_*tot*_) and MAPK ([*K*]_*tot*_), respectively. The light blue and dark blue markers in each of the panels (f-j) demarcate the portion of the trajectories that correspond to the time series shown in panels (a-e). The steady state of the system is represented by a red marker in panels (f-i). In panels (e) and (j), the system converges to a stable limit cycle. For details of parameter values for the systems shown in each of the panels see Supplementary Information.

When the signal is withdrawn, the signaling cascade can respond with large-amplitude spiking behavior in the MAPK activity before eventually relaxing to the resting state (Fig. 3). These phenomena are seen for a range of stimuli strengths and are indicative of a form of memory that can be achieved without explicit feedback or inter-pathway crosstalk. An essential condition for observing the reverberatory activity is that prior to withdrawing the applied stimulus, the system state has been driven above the low-amplitude response regime. The complex modulations seen in these figures may arise as a result of coexisting attractors. For example, in Fig. 2 (d) the system state spends considerable time in the basin of attraction of a limit cycle before approaching a stable fixed point (see Supplementary Information for details).

**FIG. 3:**
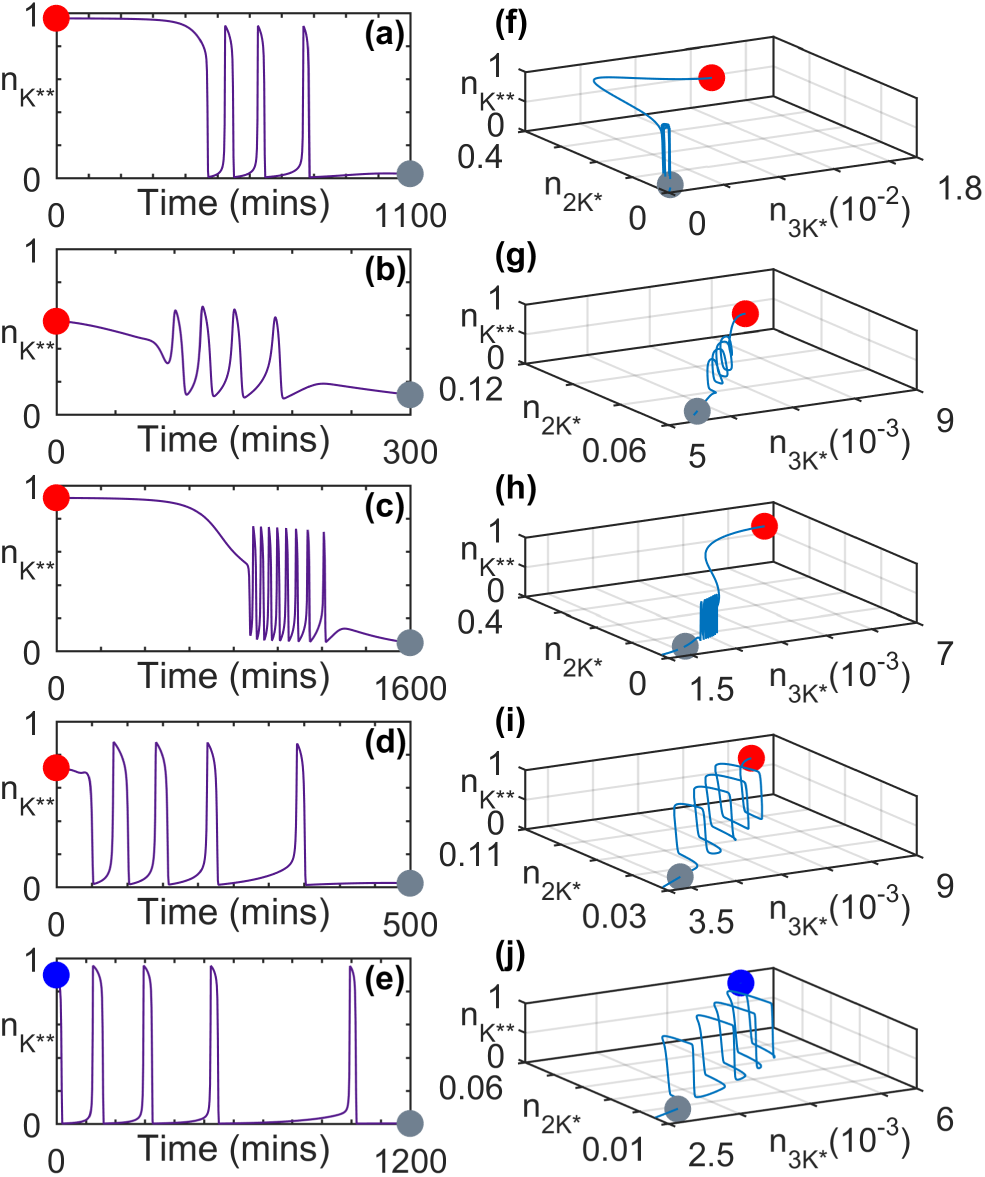
Transient activity in MAPK cascade immediately following the withdrawal (at *t* = 0) of an applied stimulus having amplitude *S* = 1.2 × 10^−6^*µM*. (a-e) Characteristic time series for the normalized concentration of doubly phosphorylated MAPK (*n_K**_*) shown for different total concentrations of kinases. (f-j) Trajectories representing the evolution of the systems in panels (a-e) in the projection of the phase-space on the planes comprising normalized concentrations of active MAP3K (*n_3K*_*), singly phosphorylated MAPK (*n_2K*_*) and active MAPK (*n_K**_*). The concentrations have been normalized by the total concentration of MAP3K ([3*K*]_*tot*_), MAP2K ([2*K*]_*tot*_) and MAPK ([*K*]_*tot*_), respectively. The steady state of the system prior to the withdrawal of the stimulus is represented by a red marker (panels f-i). The system in panels (e)and (j) is seen to relax from a state characterized by stable limit cycle oscillations (represented by the blue marker). In each trajectory shown in (f-j) the grey marker denotes the state of the system corresponding to the final time point in panels (a-e). The concentration of active MAPK is close to its resting state value following the time period shown in (a-e). The parameter values for each panel are same as those for the corresponding panels in Fig. 2.

*Processes underlying long-lived memory and reverberatory dynamics.* When the stimulus is withdrawn from the MAPK cascade, the decline in MAP Kinase activity comes about through MAPK** binding to MAPK P’ase which dephosphorylates it, resulting in an increased concentration of MAPK* [Step 1, Figs. 1(a) and 4(a)]. In turn, the phosphatase binds to MAPK* thereby deactivating it to MAPK which results in an extremely rapid decline in the concentration of MAPK* (Step 2). Concurrently, the deactivation of MAP2K** is delayed, as most of it is bound in the complex MAP2K**.MAPK that has a long time-scale of disassociation. To proceed further we can analyze the constituent processes in terms of the normalized chemical flux N_Flux_ of a molecular species, i.e., its rate of growth expressed relative to the maximum rate of growth of MAPK**. We observe that the suppression of MAP2K^**^ deactivation mentioned above results in its normalized chemical flux exceeding that of MAPK [Fig. 4(b)]. Thus, there is a net growth in activity in the MAP Kinase layer as whenever MAP2K** is released from the complex, it is available to phosphory-late MAPK which results in an increase in the concentration of MAPK^*^ (Step 3). The resulting rise in MAPK^*^ manifests as a spike in its concentration [Fig. 4(a)], and it subsequently gets phosphorylated again to increase MAPK^**^ concentration even in the absence of any stimulation (Step 4). When the net difference between the normalized flux of MAP2K^**^ and MAPK reaches a maximum, the normalized chemical flux of MAPK^**^ attains its highest value and consequently peak activity of MAP

**FIG. 4:**
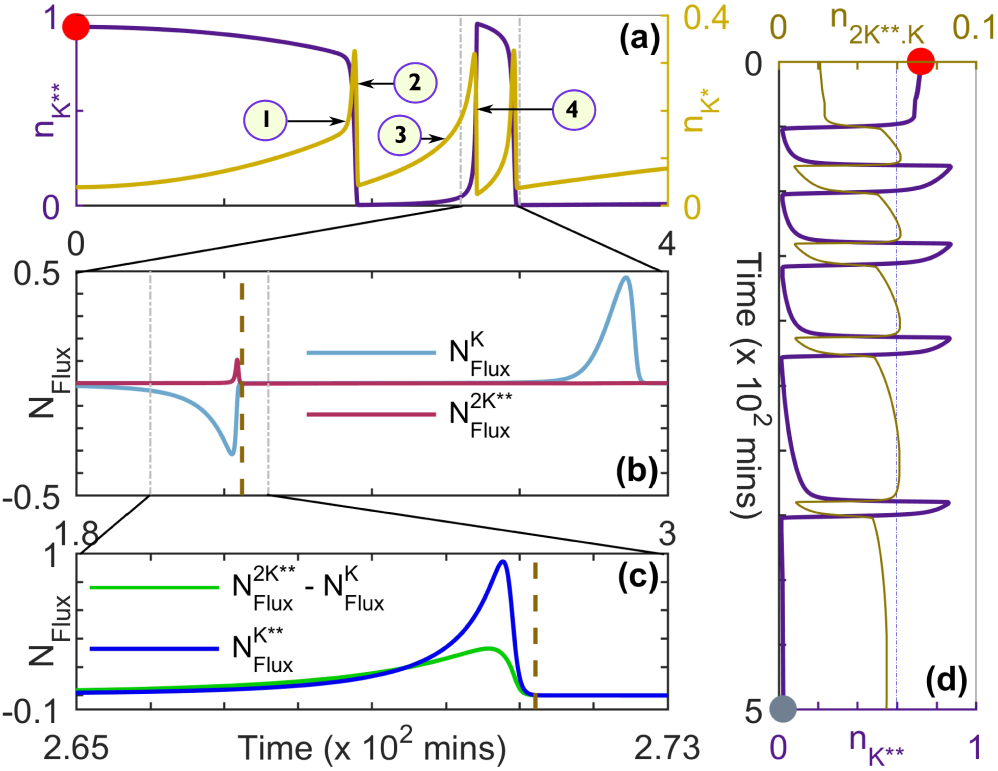
Processes underlying emergent memory and reverberatory dynamics in the MAPK cascade. (a) A characteristic time-series for the normalized concentrations of singly and doubly phosphorylated MAPK (*n_K*_* and *n_K**_*, respectively) following the removal of an applied stimulus of amplitude *S*= 2.0 × 10^−6^ *µM* at *t* = 0. The numbers (1 − 4) represent the sequence of events that lead to the emergence of the post-stimulus large-amplitude spiking activity shown schematically in Fig. 1 (b). (b) Normalized chemical flux *N*_Flux_ of MAPK and MAP2K^**^ shown for the segment of the time-series where the spiking behavior in *n_K**_* is observed following the withdrawal of the stimulus to MAP3K [demarcated by broken vertical lines in (a)]. (c) Normalized chemical flux *N*_Flux_ of MAPK^**^ shown along with the difference between the normalized fluxes of MAP2K^**^ and MAPK for the duration indicated by broken vertical lines in (b) corresponding to the peak in the spiking activity of MAPK^**^. For both panels (b) and (c), normalization of flux is with respect to the maximum of the flux for MAPK^**^. (d) Characteristic time-series for the reverberatory activity of MAPK following the withdrawal of a stimulus of amplitude *S* = 1.2 × 10^−6^ *µM* at *t* = 0, showing the normalized concentration of MAPK^**^ (*n_K**_*) along with that of the protein complex MAP2K^**^.MAPK (*n_2K**_* = [MAP2K^**^.MAPK]/[2*K*]_*tot*_). The reference line shows that the peak normalized concentration of the protein complex eventually decreases over time. For details of parameter values for (a-c) see Supplementary Information. The parameter values for panel (d) are same as those in Fig. 3(d). The steady state of the system prior to the withdrawal of the stimulus is represented by a red marker [panels (a) and (d)] while the grey marker in (d) corresponds the final time point in Fig. 3(d).

Kinase is observed [Fig. 4(c)]. Thus, steps 1-4 represent one complete cycle of MAP Kinase reverberatory activity characterized by an initial decline and a subsequent rise in MAPK^**^ concentration. These steps are subsequently repeated a number of times resulting in a series of spikes in MAPK activity [Fig. 4(d)]. The abrupt nature of the rise and fall of MAP Kinase activity that manifests as spikes is a consequence of the bistable nature of the dynamics in the MAPK layer of the cascade [15, 17]. We note that similar spiking behavior is also observed in the activity of MAP2K, with the phase of the MAP2K^**^ spikes shifted slightly forward with respect to the corresponding ones in MAPK^**^, which suggests that they result from retrograde propagation of activity from the MAPK to the MAP2K layer [14]. On the other hand, MAP3K shows a monotonic decline in its activity following the removal of the stimulus.

In order to characterize in detail the memory of prior activity retained by the cascade which is manifested as long-lived transient reverberations following the withdrawal of stimulus, we use the following measures (see Methods): (i) the primary recovery time (*τ_PR_*), (ii) the number of spikes (*N_r_*) that occur during the relaxation process, (iii) the temporal intervals between successive spikes (*t_i_* − *t*_*i*−1_, where *t_i_* is the time of occurrence of the ith spike event) and (iv) the total duration of reverberatory activity (*τ_r_*) following primary recovery. The total memory time (*τ_m_*) is the sum of *τ_PR_* and *τ_r_*as indicated in Fig. 1 (b). In the following we use these measures to present a detailed characterization of the behavior of the cascade components over a range of parameter values (Figs. 5-7).

**FIG. 5:**
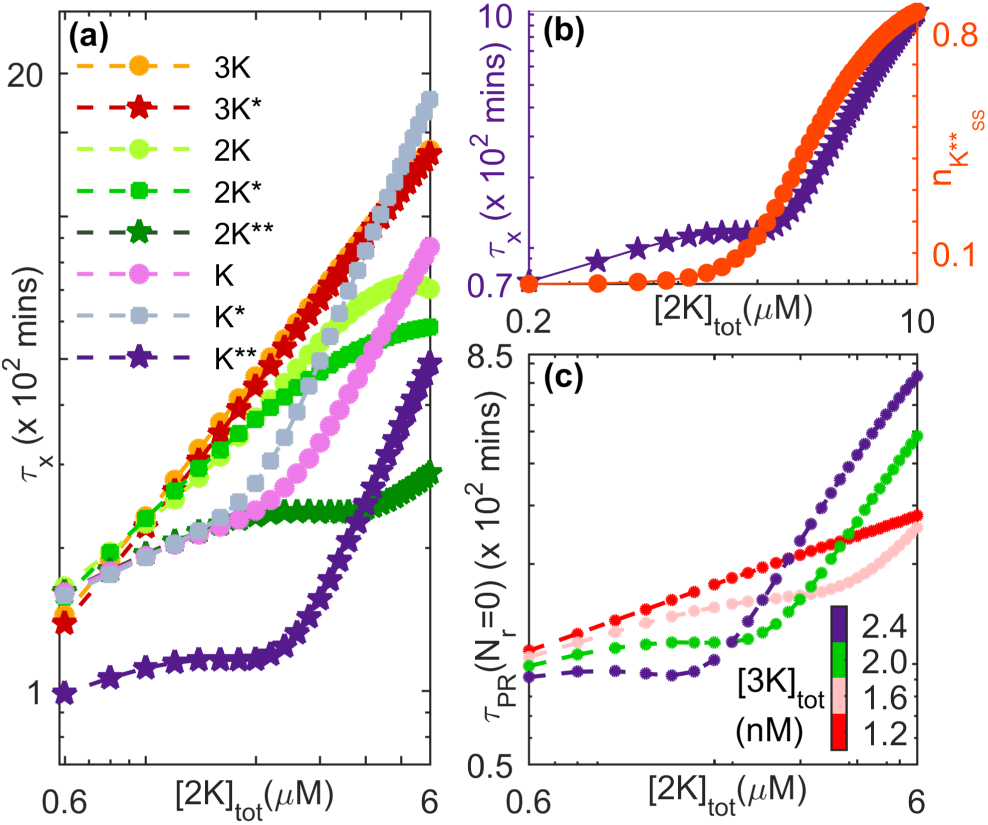
Components of the MAPK cascade exhibit relaxation behavior occurring over a broad range of time-scales. Decay of activity is shown after withdrawing an applied stimulus of amplitude *S* = 1.2 × 10^−6^*µM*. (a) The relaxation times *τ*_x_ of the different molecular species (non-, singly-and doubly phosphorylated kinase proteins) in each of the layers of the cascade vary with the total concentration of MAP2K. The nature of this dependence is distinct for lower and higher values of [2*K*]_*tot*_, which is most prominently observed in the lower layers of the cascade. (b) The occurrence of distinct regimes in the relaxation behavior of MAPK^**^ for different [2*K*]_*tot*_ is related to the corresponding increase in the steady state value attained by MAPK^**^ concentration under sus-tained stimulation of the cascade. At a specific value of the steady-state normalized MAPK activity *n_K**_*, we observe a crossover from the regime characterized by slowly increasing *τ*_x_ seen at lower total concentrations of MAP2K to a regime where *τ*_x_ increases relatively rapidly for higher [2*K*]_*tot*_. (c) The crossover behavior is also observed in the dependence of the closely related measure *τ_PR_*, the primary recovery time (see Methods), on [2*K*]_*tot*_. The difference between the two regimes become more prominent upon increasing the total concentration of MAP3K ([3*K*]_*tot*_). For both panels (a) and (b)[*K*]_*tot*_ = 0.8*µM* and [3*K*]_*tot*_ = 2.0nM, while for panel (c), [*K*]_*tot*_ = 0.8*µM*. For details of all other parameter values see Supplementary Information.

**FIG. 6:**
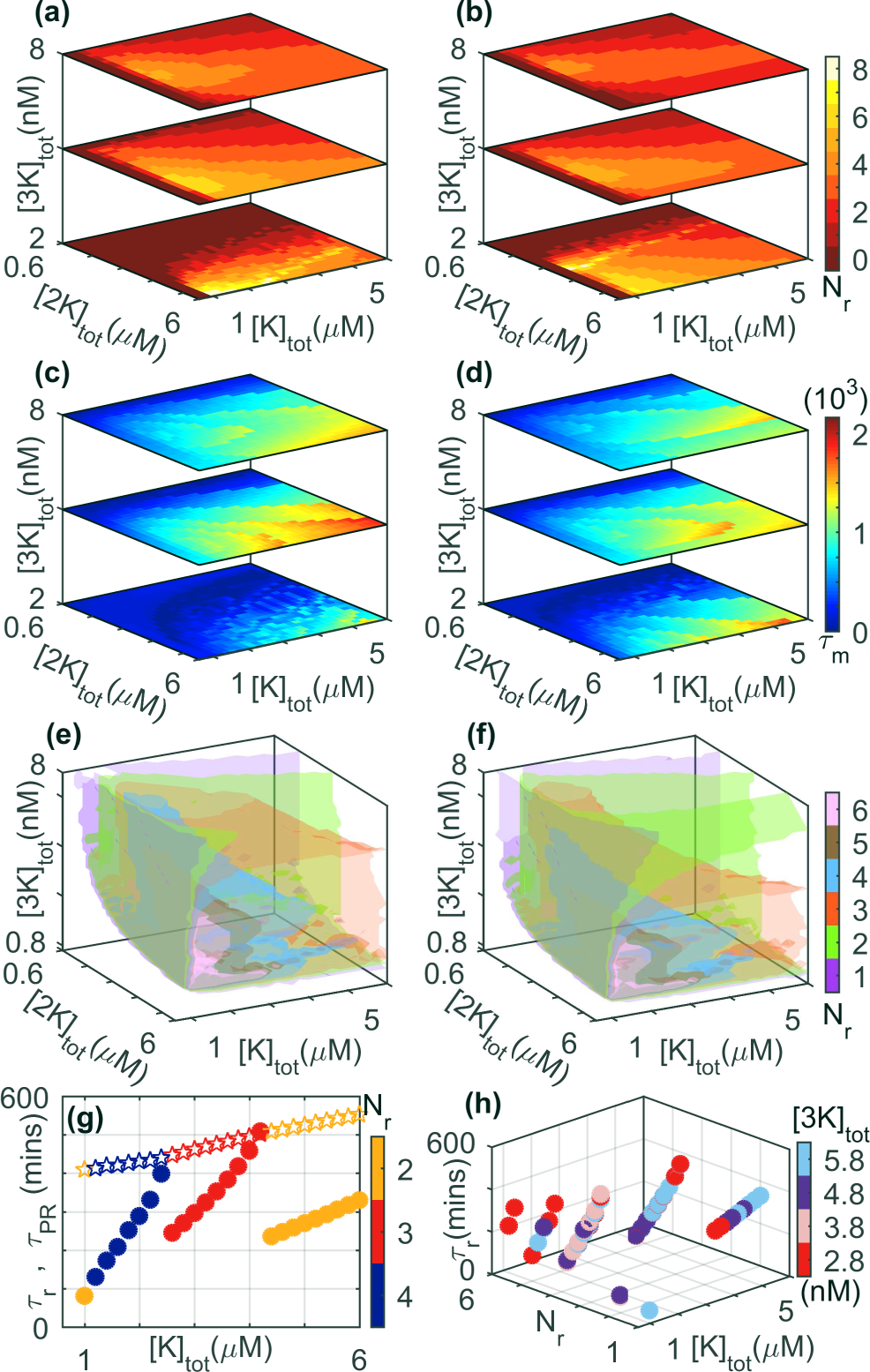
Dependence of reverberatory activity on the total kinase concentrations, viz., [*K*]_*tot*_, [2*K*]_*tot*_ and [3*K*]_*tot*_. (a-b) The number of spikes *N_r_*, (c-d) the total memory time *τ_m_* (in minutes) and (e-f) isosurfaces for *N_r_* observed on withdrawing an applied stimulus of amplitude *S* [= 0.8 × 10^−6^ *µM* for (a,c,e) and 1.2 × 10^−6^*µM* for (b,d,f)] are shown as functions of total concentrations of the three kinases. (g) The primary recovery time *τ_PR_* (stars) and the total duration of reverberatory activity *τ_r_*(filled circles) are shown for different values of *N_r_* (indicated by the color bar). While *τ_PR_* increases monotonically with increasing total MAPK concentration, *τ_r_*shows a more complex dependence ([2*K*]_*tot*_ = 3*µM* and [3*K*]_*tot*_ = 4nM). (h) The dependence of *τ_r_*on [*K*]_*tot*_ for different values of *N_r_* has a similar nature for different choices of [3*K*]_*tot*_ (indicated by the color bar, [2*K*]_*tot*_ = 3*µM*). Note that for panel (h), we consider only situations where the system attains a steady state on maintaining stimulation. For details of all other parameter values see Supplementary Information.

*MAP Kinase cascade components have different recovery timescales.* As mentioned earlier, the emergence of long-lived reverberatory activity of MAPK following the withdrawal of an applied stimulus can be linked to the flux imbalance of different cascade components which suggests significant differences in their rates of relaxation. As shown in Fig. 5(a), this is indeed the case, even for parameter regimes where no spiking activity of MAPK is observed (i.e., *N_r_* = 0). As can be seen, the nature of increase of the relaxation time with increasing total concentrations of kinase protein MAP2K is distinct for the different molecular species and also depends on the state of their phosphorylation. In the lower layers of the cascade, we also find a crossover between two regimes seen at lower and higher values of [2*K*]_*tot*_ respectively. These regimes are characterized by relatively slow and rapid increases (respectively) in the recovery times with increasing [2*K*]_*tot*_, and appear to be related to the steady-state value attained by MAPK activity upon sustained stimulation of the cascade for the corresponding value of [2*K*]_*tot*_ [Fig. 5 (b)]. The crossover between the two regimes is seen to occur for a value of [2*K*]_*tot*_ for which ~ 17% of MAPK is activated for the parameter values used in Fig. 5(b).

The distinct regimes are also observed in the dependence of the primary recovery time *τ_PR_* on [2*K*]_*tot*_ [Fig. 5(c)]. As can be observed, the difference between the regimes becomes more pronounced with an increase in the total concentration of MAP3K. An important point to note is that for lower values of [2*K*]_*tot*_, the recovery time decreases with increasing [3*K*]_*tot*_ while the reverse trend is seen for higher values of [2*K*]_*tot*_. We have verified that increasing the stimulus amplitude S while keeping the total MAP3K concentration fixed has a similar effect on the relaxation behavior of activated MAPK (see Supplementary Information). As increasing total concentration of MAP2K results in increased steady-state activity of MAPK, we conclude that, in general, higher activity states of MAPK are associated with increasing relaxation time when either the signal or the substrate (MAP3K) is increased. Conversely, for states characterized by much lower MAPK activity, larger values of *S* or [3*K*]_*tot*_ results in reduced relaxation periods.

*Dependence of reverberatory activity on total kinase concentrations.* Diverse cellular environments are characterized by different total concentrations of the various molecular components of the MAPK cascade. Thus, in order to determine the robustness of spiking and reverberatory activity following the removal of an applied stimulus, it is important to see how they are affected by varying total kinase concentrations. Such a study will also indicate the ease with which these phenomena can be experimentally observed. Fig. 6 shows the variation of different measures of reverberatory activity on the total concentrations of MAPK, MAP2K and MAP3K. While there is a complex dependence on these parameters for the exact number of spikes *N_r_* and the duration of the total memory time *τ_m_*, the phenomenon of reverberatory activity following withdrawal of stimulation can be observed over a large range of the parameter space, underlining its robustness. We also observe that on increasing [3*K*]_*tot*_, the response of *N_r_* to variation in [*K*]_*tot*_ and [2*K*]_*tot*_ becomes relatively homogeneous. Increasing the stimulus amplitude S [compare panels (a,c,e) with (b,d,f) of Fig. 6] does not seem to alter the qualitative nature of the variation in *N_r_* and *τ_m_* over the parameter space in general, although we do observe that the domains corresponding to different values of *N_r_* occupy different regions [Fig. 6(e and f)]. Note that for low [3*K*]_*tot*_, high values of *N_r_* are observed to coexist with low values of τm [Fig. 6(a,c and b,d)]. While it may appear surprising that these two measures of memory are not in consonance in this region of parameter space, it can be explained by noting that the stimulated system is in an oscillatory state, and following the removal of the signal these relatively high-frequency oscillations cease after a short duration. Fig. 6(g) suggests that the variation seen in *τ_m_* as a function of the total MAPK concentration for a specific *N_r_* is mostly governed by *τ_r_*, the total duration of reverberatory activity, with the corresponding dependence of *τ_PR_* on [*K*]_*tot*_ being weak.

As the total MAPK concentration is increased, we observe that while the primary recovery time increases almost linearly, the nature of the reverberatory dynamics as reflected in *τ_r_* shows a more complex dependence on [*K*]_*tot*_ [Fig. 6(g)]. If for a given value of [*K*]*_tot_* the MAPK activity following withdrawal of the stimulus shows *N_r_* spikes over a duration of *τ_r_*, then on increasing [*K*]_*tot*_ the time-interval between the spikes increases (thereby resulting in an increase of *τ_r_*) until a critical value beyond which the last of the *N_r_* spike no longer appears. Thus, at this point *N_r_* reduces by unity with a concomitant drop in *τ_r_*. This series of events is repeated for steadily decreasing values of *N_r_* as the total MAPK concentration is increased further. Each value of *N_r_* is associated with a characteristic rate of increase in *τ_r_*with [*K*]_*tot*_. With a reduction in *N_r_* (as a result of increasing [*K*]_*tot*_), this rate is found to decrease as well, which suggests a saturation of the system response. These results are robust with respect to different choices of total MAP3K concentration as can be seen from Fig. 6(h), suggesting that similar behavior will be seen for a range of strengths for the applied signal (see Supplementary Information).

*Dependence of reverberatory activity on total phosphatase concentrations.* We have also investigated the role that phosphatase availability plays on the reverberatory activity of the cascade following the withdrawal of the stimulus. As is the case for total kinase concentrations shown in Fig. 6, we see from Fig. 7 (a-b) that the number of spikes *N_r_* and the duration of total memory time *τ_m_* depend on the total concentrations of the phosphatases MAPK P’ase, MAP2K P’ase, and MAP3K P’ase. For larger values of the concentrations, viz., [*P_K_*], [*P_2K_*] and [*P_3K_*], respectively, the system operates in the low-amplitude response regime. As mentioned earlier,the reverberatory MAPK dynamics during recovery following withdrawal of the applied stimulus will not be seen in this regime. As the phosphatase concentrations are decreased, spiking behavior of MAPK activity is observed with both *τ_m_* and *N_r_* attaining high values in an optimal range. The large variation seen in *τ_m_* [Fig. 7 (b)] arises as regions in [*P_2K_*]-[*P_3K_*] parameter space characterized by the same value of *N_r_* are seen to exhibit a range of different values of *τ_r_*and τ_P R_ [Fig. 7 (d)]. For reverberatory activity associated with a specific *N_r_*, we observe that the duration *τ_r_*increases with increasing total MAPK P’ase concentration. This is a consequence of the intervals between successive spikes (*t_i_* − *t*_*i*−1_) increasing with [*P_K_*] as is shown in Fig. 7 (c). Note that the results are qualitatively similar for different amplitudes of the applied stimulus (see Supplementary Information). However, increasing [*P_K_*] results also in decreased time for primary recovery *τ_PR_* [Fig. 7 (d)], which in conjunction with the previously mentioned result leads to non-monotonic dependence of the total memory time *τ_m_* on phosphatase availability. While this non-monotonicity is suggested in Fig. 7 (b), it is shown clearly in Fig. 7 (e) where the central panel corresponds to situations where spiking behavior is observed in MAPK activity. Investigation into the dependence of *τ_m_* on *P_K_*[Fig. 7(e)] reveals that the range of [*P_K_*] over which reverberatory activity (i.e., *N_r_* =6 0) occurs is demarcated by discontinuities in the functional dependence of *τ_m_* on *P_K_*. For intermediate *P_2K_*[Fig. 7(e), central panel] where the system attains a steady state on maintaining stimulation, the spiking activity following withdrawal of the stimulus becomes more prominent for low total concentration of MAP3K P’ase. For higher *P_2K_*[Fig. 7(e), right panel] where the system becomes oscillatory over an intermediate range of [*P_K_*], reverberatory activity is observed over a broader range of [*P_3K_*]. While we have assumed that the same phosphatase acts on both the singly and doubly phosphorylated forms of the kinase in a particular layer of the cascade (as in the canonical Huang-Ferrell model), we have explicitly verified that our results are not sensitively dependent on this.

**FIG. 7:**
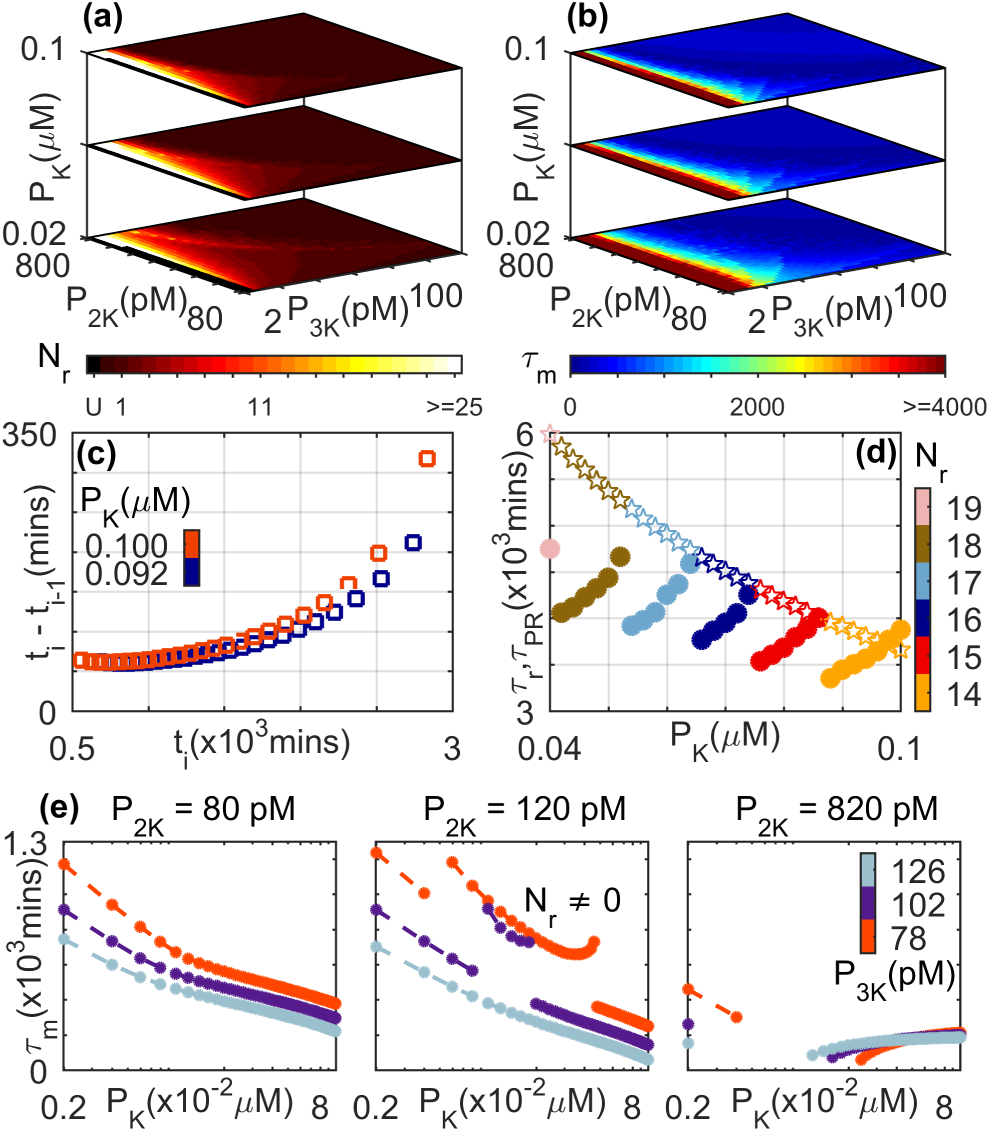
Dependence of reverberatory activity on the total concentrations of the phosphatases MAPK P’ase ([*P_K_*]), MAP2K P’ase, ([*P_2K_*]) and MAP3K P’ase ([*P_3K_*]). (a) The number of spikes *N_r_* and (b) the total memory time *τ_m_* (in minutes) observed on withdrawing an applied stimulus of amplitude S = 0.8 × 10^−6^ *µM*. Situations where the primary recovery time is longer than a maximum or cut-off value (see Methods), such that the duration of the reverberatory dynamics cannot be properly measured, are indicated by the color corresponding to “U”. (c) The interval between successive spikes *i* − 1 and i increases with time (*t_i_* being the time of occurrence of the ith spike). As the MAPK P’ase concentration is increased, the durations of these intervals are seen to increase. The total concentrations of the other two phosphatases are maintained at [*P_2K_*] = 680pM and [*P_3K_*] = 10pM. (d) The variation of primary recovery time *τ_PR_* (stars) and the total duration of reverberatory activity *τ_r_*(filled circles) as a function of total MAPK P’ase concentration are shown for different values of *N_r_* (indicated by the color bar). While *τ_PR_* decreases monotonically with increasing [*P_K_*], *τ_r_*shows a more complex dependence ([P_2K_] = 200pM and [*P_3K_*] = 6pM). (e) Dependence of the total memory time *τ_m_* on total MAPK P’ase concentration ([*P_K_*] shown in log scale) for different total concentrations of MAP2K P’ase (values indicated above each of the three panels) and MAP3K P’ase (indicated using different colors as shown in the color bar). Note that we consider only situations where the system attains a steady state on maintaining stimulation. For details of all other parameter values see Supplementary Information.

## IV. DISCUSSION

In this paper we have shown that an isolated MAPK signaling module can serve as a fundamental motif in the intra-cellular signaling network for imparting a form of short-term memory to the cell. The emergence of long-lived reverberatory activity reported here arises from the diversity of relaxation timescales for the different components of the MAP Kinase cascade, which results in flux imbalance between activation of the MAPK layer and deactivation in the MAP2K layer. One may therefore expect to observe results qualitatively similar to what has been reported here whenever the system has disparate timescales regardless of the actual molecular concentrations and kinetic rates which can vary substantially across different cells [32–34]. Thus, as the MAPK cascade is present in all eukaryotic cells [5, 6], the mechanism for short-term memory in such a signaling cascade that is presented here may hold for such cells in general. As the duration of MAPK^**^ activity is critical for many cellular decisions [35], e.g., the prolonged activation of ERK resulting in its translocation to the nucleus [36], the persistent reverberatory activity seen here may play a non-trivial role in regulation of cellular functions.

The basal level activity of MAPK in a normal cell is maintained at a low proportion of the total MAPK concentration and serves several biological functions [37]. We observe a crossover between two qualitatively distinct regimes of relaxation behavior of MAPK^**^ occurring at a steady state that is characterized by relatively low proportion of activation of the available MAPK [~ 17% in Fig. 5 (b)]. Thus, there appears to be an effective threshold for MAPK activity (which may be related to its basal state level) that demarcates the different relaxation regimes following the removal of the applied stimulus. A similar crossover is also observed for the primary recovery time *τ_PR_*.

It is known that ERK MAPK isoforms (e.g., p42 and p44) are abundantly expressed in non-dividing terminally differentiated neurons [27]. Activation of MAPK by spaced stimulation is known to be responsible for morphological changes in dendrites [26]. Studies also suggest that the activation of the MAPK pathway is linked with associative learning in the mammalian nervous system, synaptic plasticity and neurological memory [25–27, 38, 39]. An intriguing possibility suggested by the results reported here is that the observed repeated spiking in MAPK activity may function as an effective temporally spaced signal to the nucleus of a neuron. This can then facilitate subsequent changes in the cell required for memory formation.

Another well-known example of eukaryotic cellular memory is observed during chemotactic migration along the gradient of a chemical signal [40, 41]. The directionality of migration is known to persist for a certain duration, even if the chemical gradient is altered or becomes static. Studies show that the protein Moesin contributes to the long-lived rigidity of the cytoskeleton assembly that subsequently leads to the directional memory in polarized migrating cells [41]. However, the intra-cellular processes that underlie the persistent activity of Moesin in the absence of a gradient mediated signal are still largely unknown. Evidence suggests that the regulation of Moesin and other ERM proteins are linked with the activity of the MAPK pathway [42, 43]. The long-term reverberatory activity of MAPK following the withdrawal of a stimulus that is reported here may be a possible mechanism underlying such persistent cellular behavior.

To conclude, we have shown the possibility of long-lived reverberatory activity in a signaling cascade following the withdrawal of external stimuli. Our results suggest a mechanism through which the intra-cellular signaling system can encode short-term memory of signals to which the cell was previously exposed. The large-amplitude spiking activity of MAPK following the removal of a prior stimulus may also provide a mechanism for signal integration and learning when the cascade is repeatedly stimulated. We note that there may be additional factors not considered here that may lengthen the persistence of reverberatory activity, including scaffold proteins that increase the lifetime of kinase complexes. Our results suggest that the MAPK cascade potentially has a key role in shaping the information processing capabilities of eukaryotic cells in diverse environments.

## Acknowledgments

SNM is supported by the IMSc Complex Systems Project (12^th^ Plan). The simulations required for this work were done in the High Performance Computing facility (Nandadevi and Satpura) of IMSc which is partially funded by DST (SR/NM/NS-44/2009). We thank James Ferrell, Upinder Bhalla, Tharmaraj Jesan, Uddipan Sarma, Bhaskar Saha, Jose Faro, Vineeta Bal, J. Krishnan, Mukund Thattai, Marsha Rosner and Pamela Silver for helpful discussions.

## SUPPLEMENTARY INFORMATION

### S1. THE MODEL EQUATIONS

**TABLE S1:** Components of the MAPK Cascade

| Component | Notation | Symbol |
| --- | --- | --- |
| Mitogen-activated Protein Kinase Kinase Kinase | MAP3K | 3K |
| Singly Phosphorylated Mitogen-activated Protein Kinase Kinase Kinase | MAP3K* | 3K* |
| Mitogen-activated Protein Kinase Kinase | MAP2K | 2K |
| Singly Phosphorylated Mitogen-activated Protein Kinase Kinase | MAP2K* | 2K* |
| Doubly Phosphorylated Mitogen-activated Protein Kinase Kinase | MAP2K** | 2K** |
| Mitogen-activated Protein Kinase | MAPK | K |
| Singly Phosphorylated Mitogen-activated Protein Kinase | MAPK* | K* |
| Doubly Phosphorylated Mitogen-activated Protein Kinase | MAPK** | K** |
| MAP3K-Phosphatase | 3K P'ase | P<sub>3K</sub> |
| MAP2K-Phosphatase | 2K P'ase | P<sub>2K</sub> |
| MAPK-Phosphatase | K P'ase | P<sub>K</sub> |

The three layer MAPK cascade comprises the following enzyme-substrate reactions:

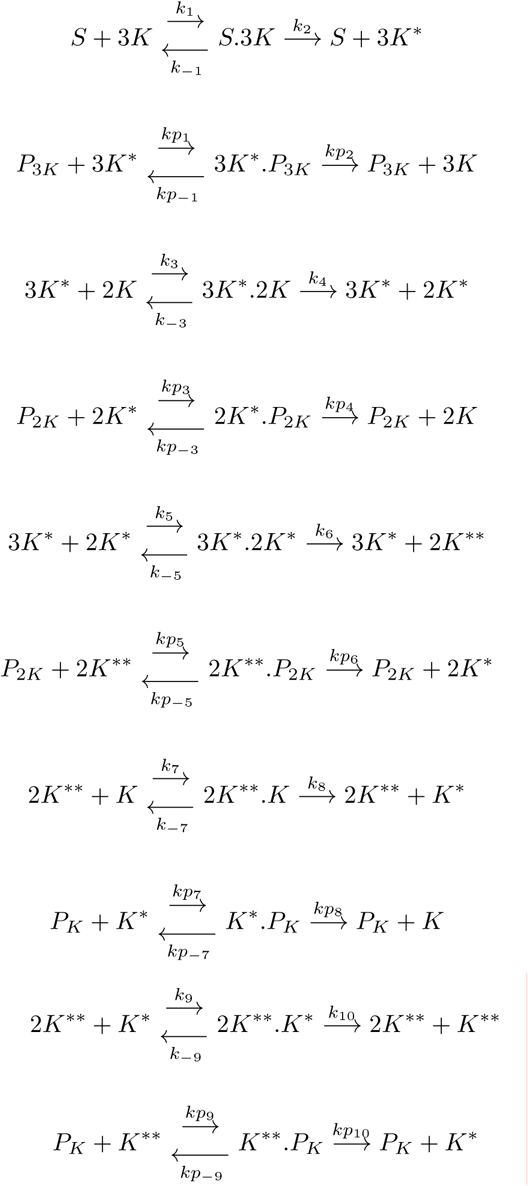

The above enzyme-substrate reactions can be expressed in terms of the following coupled ordinary differential equations (ODEs):

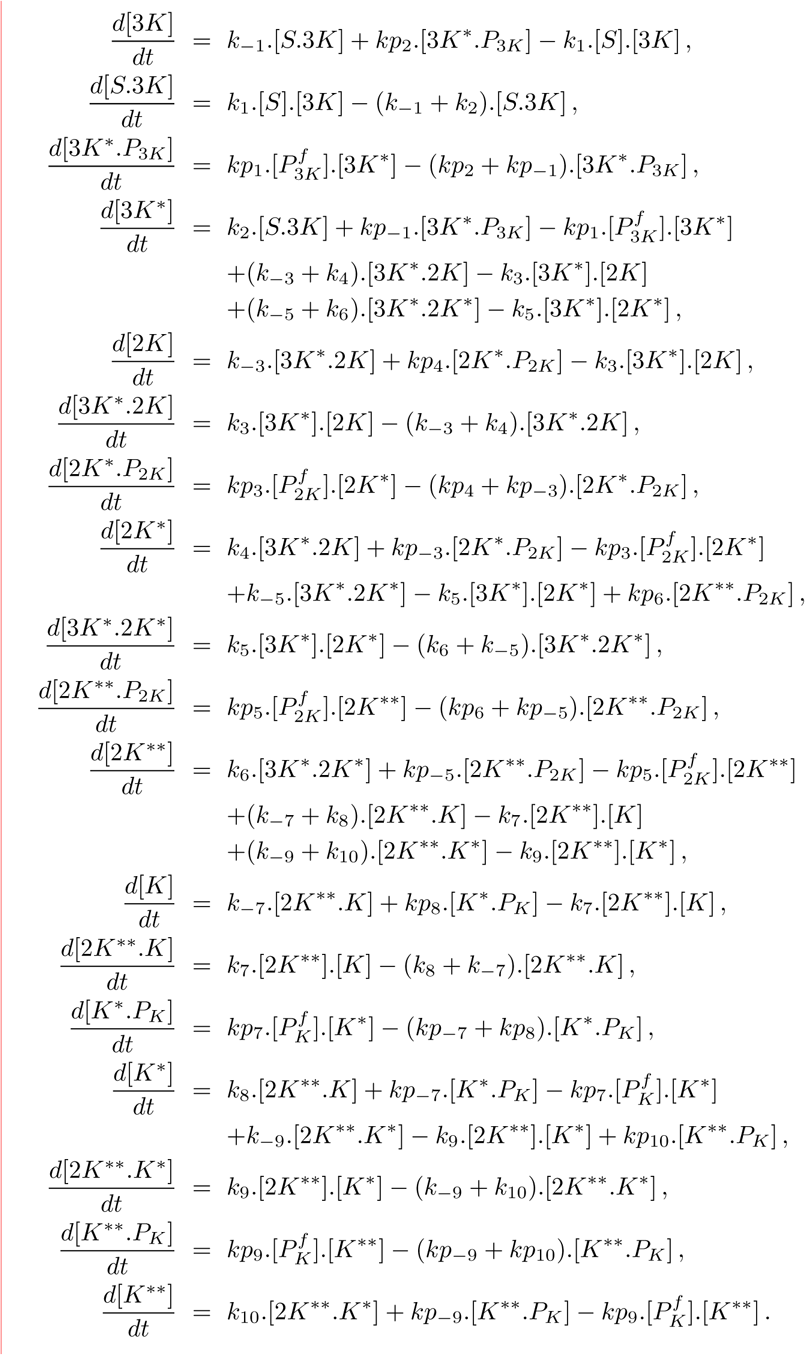

where

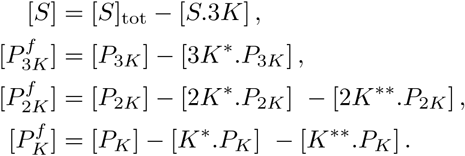

It is explicitly ensured that the total concentrations of all individual kinases and phosphatases are conserved at all times. The concentrations of the different molecular species can vary over several orders of magnitudes. We have therefore numerically solved the equations using low relative and absolute tolerances in order to ensure the accuracy of the resulting time-series.

### S2. SYSTEM PARAMETERS

The numerical values for the reaction rates are obtained from Ref. [13], and are listed in Table S2.

**TABLE S2:**
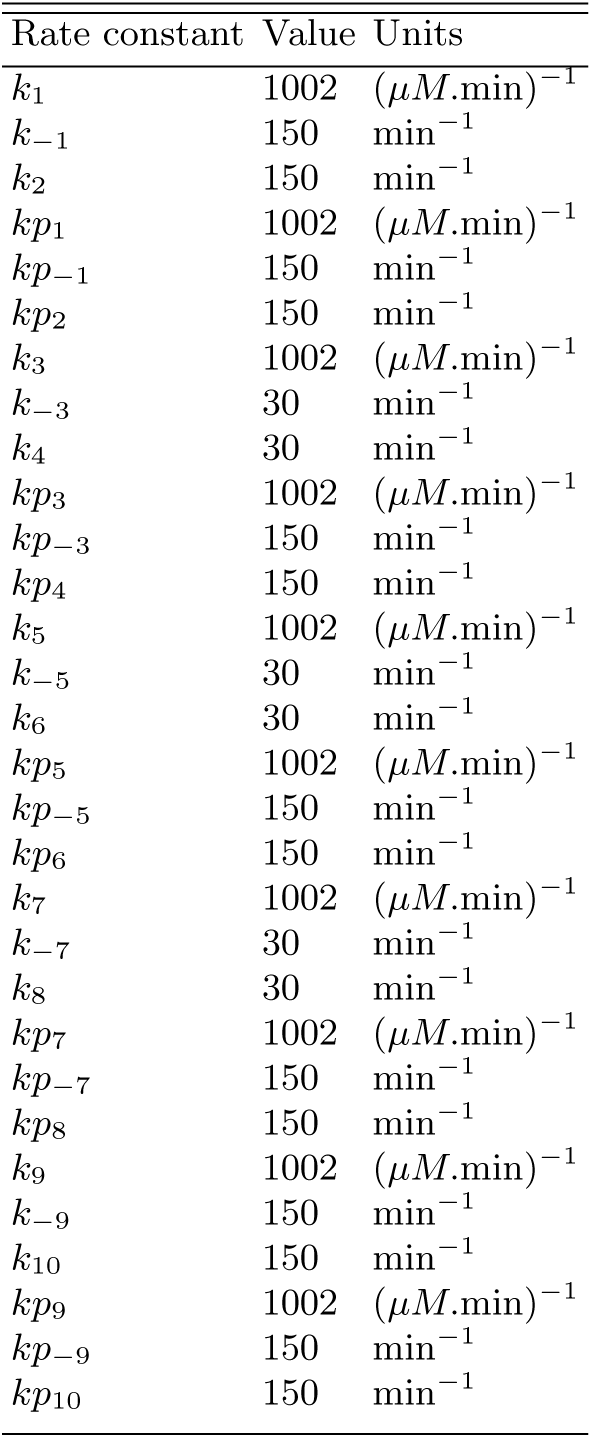
Reaction Rates

| Rate constant | Value | Units |
| --- | --- | --- |
| $k_1$ | 1002 | $(\mu M.\text{min})^{-1}$ |
| $k_{-1}$ | 150 | $\text{min}^{-1}$ |
| $k_2$ | 150 | $\text{min}^{-1}$ |
| $kp_1$ | 1002 | $(\mu M.\text{min})^{-1}$ |
| $kp_{-1}$ | 150 | $\text{min}^{-1}$ |
| $kp_2$ | 150 | $\text{min}^{-1}$ |
| $k_3$ | 1002 | $(\mu M.\text{min})^{-1}$ |
| $k_{-3}$ | 30 | $\text{min}^{-1}$ |
| $k_4$ | 30 | $\text{min}^{-1}$ |
| $kp_3$ | 1002 | $(\mu M.\text{min})^{-1}$ |
| $kp_{-3}$ | 150 | $\text{min}^{-1}$ |
| $kp_4$ | 150 | $\text{min}^{-1}$ |
| $k_5$ | 1002 | $(\mu M.\text{min})^{-1}$ |
| $k_{-5}$ | 30 | $\text{min}^{-1}$ |
| $k_6$ | 30 | $\text{min}^{-1}$ |
| $kp_5$ | 1002 | $(\mu M.\text{min})^{-1}$ |
| $kp_{-5}$ | 150 | $\text{min}^{-1}$ |
| $kp_6$ | 150 | $\text{min}^{-1}$ |
| $k_7$ | 1002 | $(\mu M.\text{min})^{-1}$ |
| $k_{-7}$ | 30 | $\text{min}^{-1}$ |
| $k_8$ | 30 | $\text{min}^{-1}$ |
| $kp_7$ | 1002 | $(\mu M.\text{min})^{-1}$ |
| $kp_{-7}$ | 150 | $\text{min}^{-1}$ |
| $kp_8$ | 150 | $\text{min}^{-1}$ |
| $k_9$ | 1002 | $(\mu M.\text{min})^{-1}$ |
| $k_{-9}$ | 150 | $\text{min}^{-1}$ |
| $k_{10}$ | 150 | $\text{min}^{-1}$ |
| $kp_9$ | 1002 | $(\mu M.\text{min})^{-1}$ |
| $kp_{-9}$ | 150 | $\text{min}^{-1}$ |
| $kp_{10}$ | 150 | $\text{min}^{-1}$ |

**TABLE S3:** Total concentration (in *µM*) of the kinase proteins for Fig. 4; (panels a–c) and Fig. 7

| $[K]_{tot}$ | $[2K]_{tot}$ | $[3K]_{tot}$ |
| --- | --- | --- |
| 4.8 | 1.2 | 0.0030 |

**TABLE S4:**
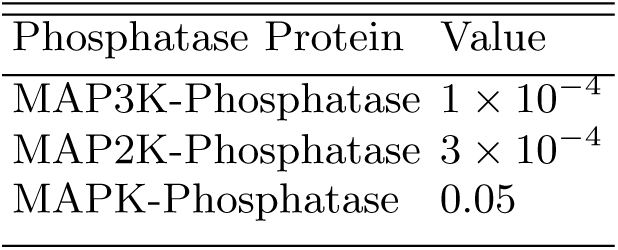
Total concentration (in *µM*) of the phosphatase proteins for Figs. 2–6

| Phosphatase Protein | Value |
| --- | --- |
| MAP3K-Phosphatase | $1 \times 10^{-4}$ |
| MAP2K-Phosphatase | $3 \times 10^{-4}$ |
| MAPK-Phosphatase | 0.05 |

**TABLE S5:**
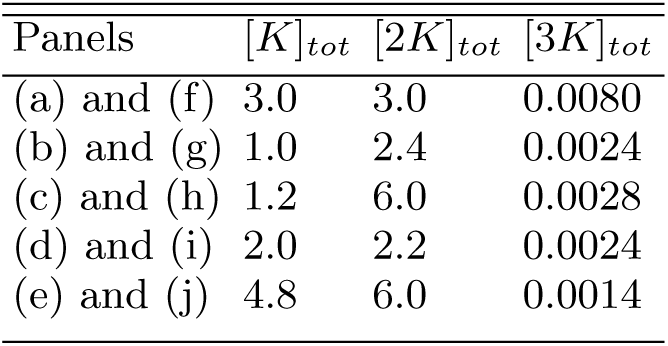
Total concentration (in *µM*) of the kinase proteins for Figs. 2–3

| Panels | $[K]_{tot}$ | $[2K]_{tot}$ | $[3K]_{tot}$ |
| --- | --- | --- | --- |
| (a) and (f) | 3.0 | 3.0 | 0.0080 |
| (b) and (g) | 1.0 | 2.4 | 0.0024 |
| (c) and (h) | 1.2 | 6.0 | 0.0028 |
| (d) and (i) | 2.0 | 2.2 | 0.0024 |
| (e) and (j) | 4.8 | 6.0 | 0.0014 |

### S3. SUPPLEMENTARY FIGURES

**FIG. S1:**
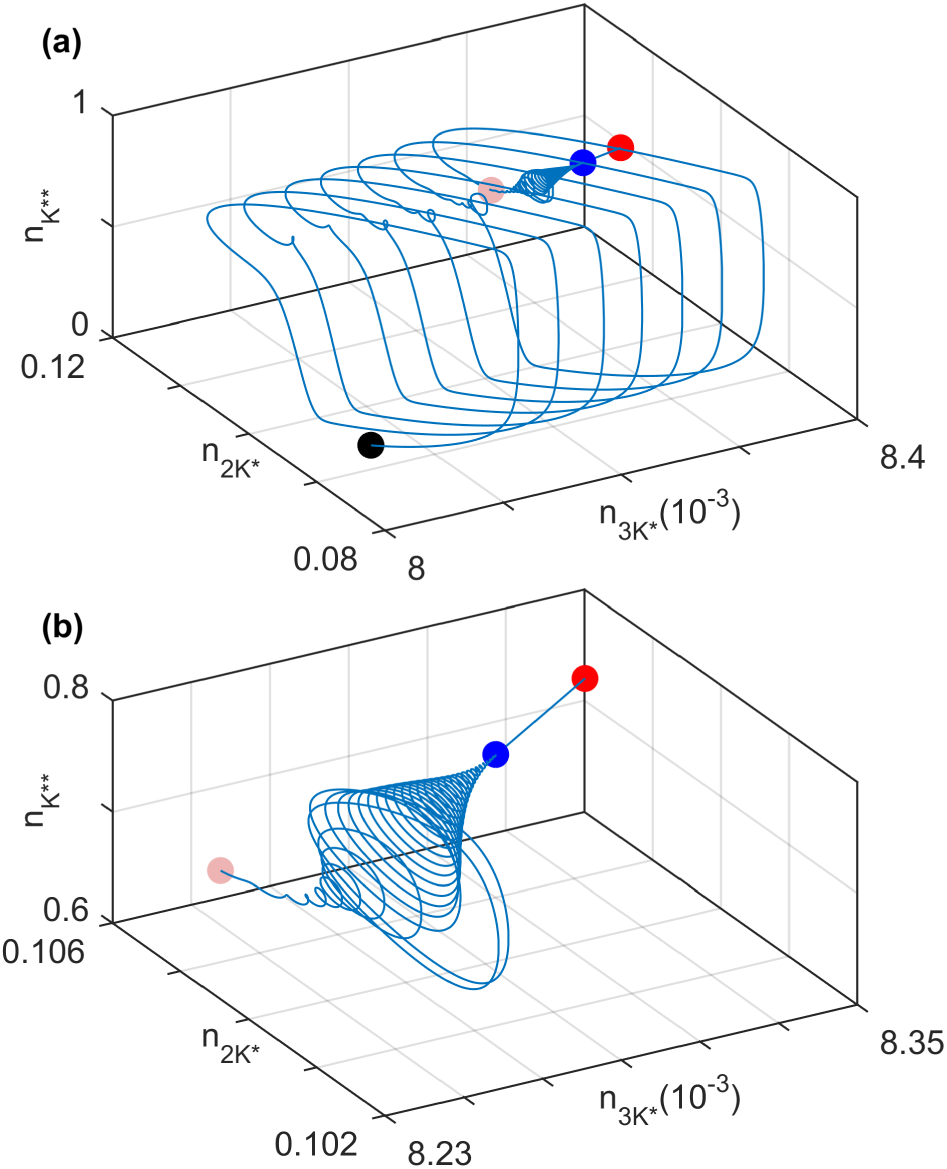
Magnified views of the phase space trajectory shown in Fig. 2(i). The blue markers correspond to the final point in the time series displayed in Fig. 2(d), while the red markers indicate the fixed point of the dynamical system in the presence of stimulus. (a) Magnified view of the trajectory beginning from the black marker shown in Fig. 2(i). The pink marker denotes the starting point of the segment of the trajectory displayed in panel (b). (b) Further magnification of the section of phase-plane trajectory shown in (a), corresponding to the duration when the system moves away from the unstable limit cycle and converges to the stable fixed point.

**FIG. S2:**
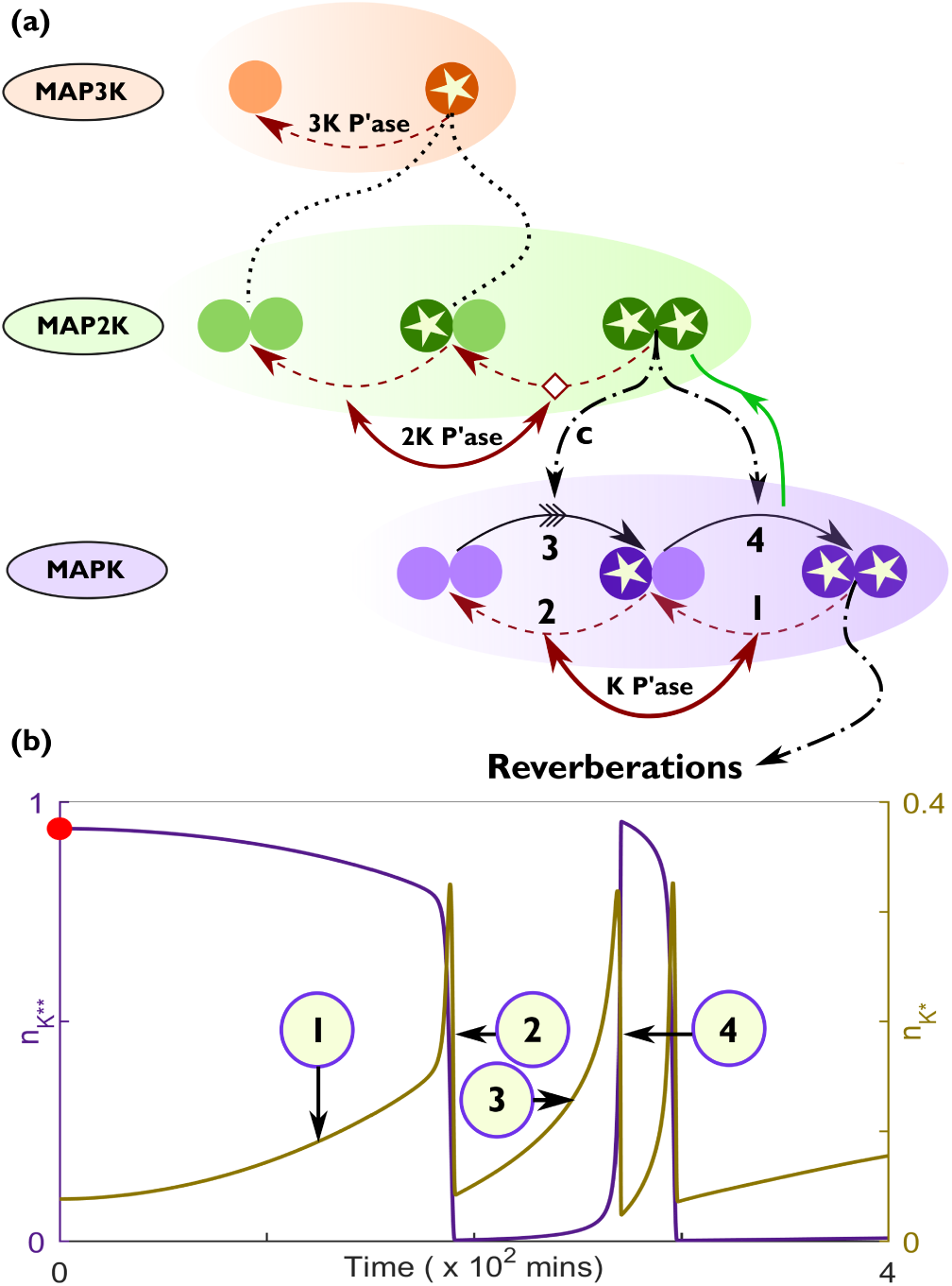
Processes underlying long-lived memory and reverberatory dynamics. (a) Schematic representation of MAPK cascade showing the processes that occur subsequent to removing a stimulus. The numbers (1 − 4) represent the sequence of events that lead to the emergence of the post-stimulus large-amplitude spiking activity shown in (b). The enzyme-substrate protein complex formed during activation of MAPK by doubly phosphorylated MAP2K is indicated by “c”. The green arrow from the MAPK layer to the MAP2K layer represents the release of doubly phosphorylated MAP2K from downstream complexes. (b) A characteristic time-series for the normalized concentration of singly and doubly phosphorylated MAPK (*n_K*_* and *n_K**_*, respectively) following the removal of an applied stimulus of amplitude *S* = 2.0 × 10^−6^ *µM* at *t* = 0. The numbers (1 − 4) represent the same events shown in (a). The total concentrations of the kinases and phosphatases used for generating the time-series are provided in Tables S3 and S4, respectively.

**FIG. S3:**
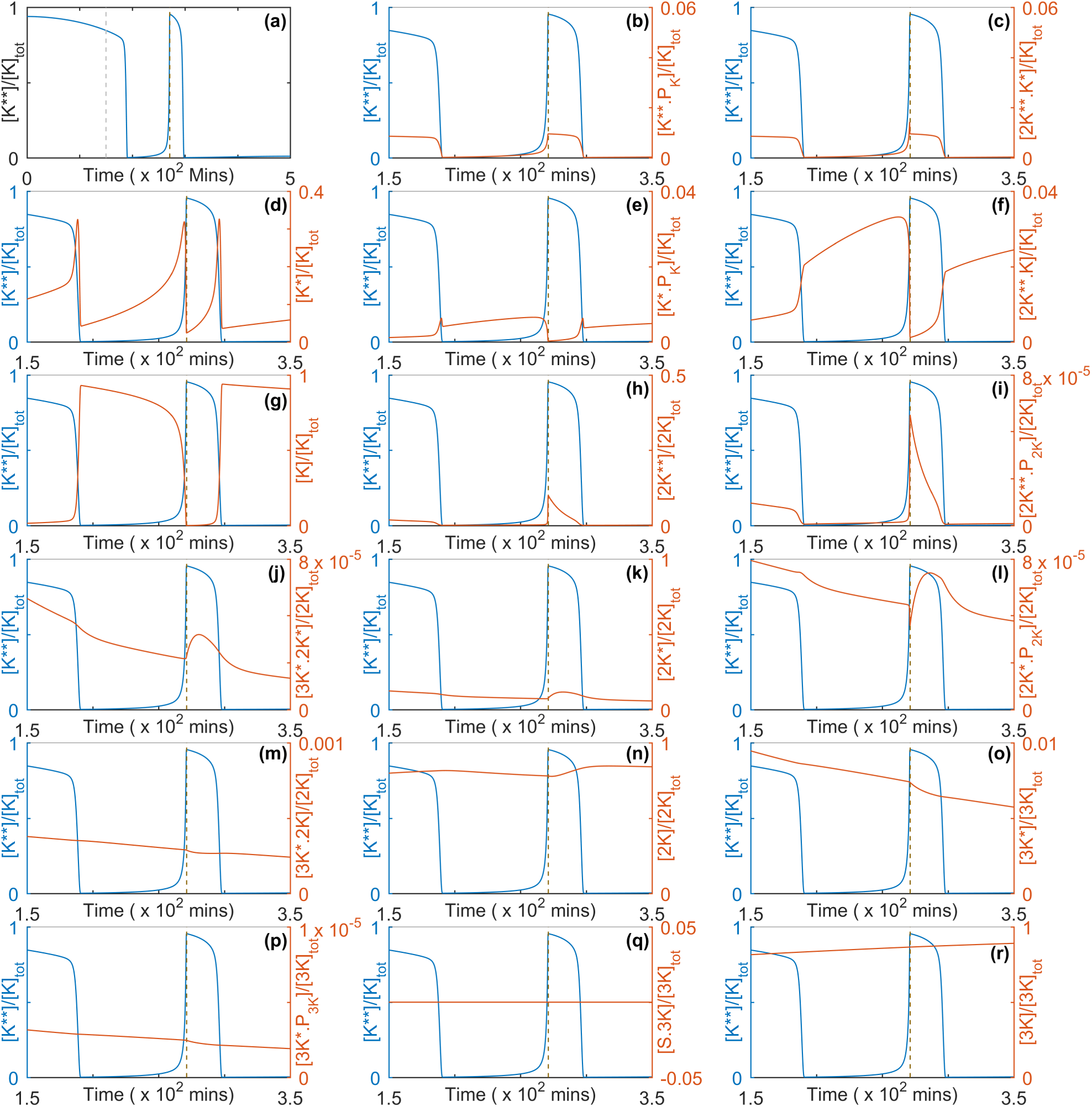
Characteristic dynamics of the molecular components of the MAP Kinase cascade following withdrawal of a stimulus. (a)The time-series of the normalized concentration of doubly phosphorylated MAPK ([*K*^**^]/[*K*]_*tot*_) following removal of an applied stimulus with amplitude S = 2.0 × 10^−6^ *µM* at t = 0. (b-r) Time-series of the normalized concentrations of the different components of the MAPK cascade, shown starting from t = 150 minutes after withdrawing the stimulus, displayed together with the time-series of normalized MAPK activity [*K*^**^]/[*K*]_*tot*_. The total concentrations of the kinases and phosphatases used for generating the figures are provided in Tables S3 and S4, respectively.

**FIG. S4:**
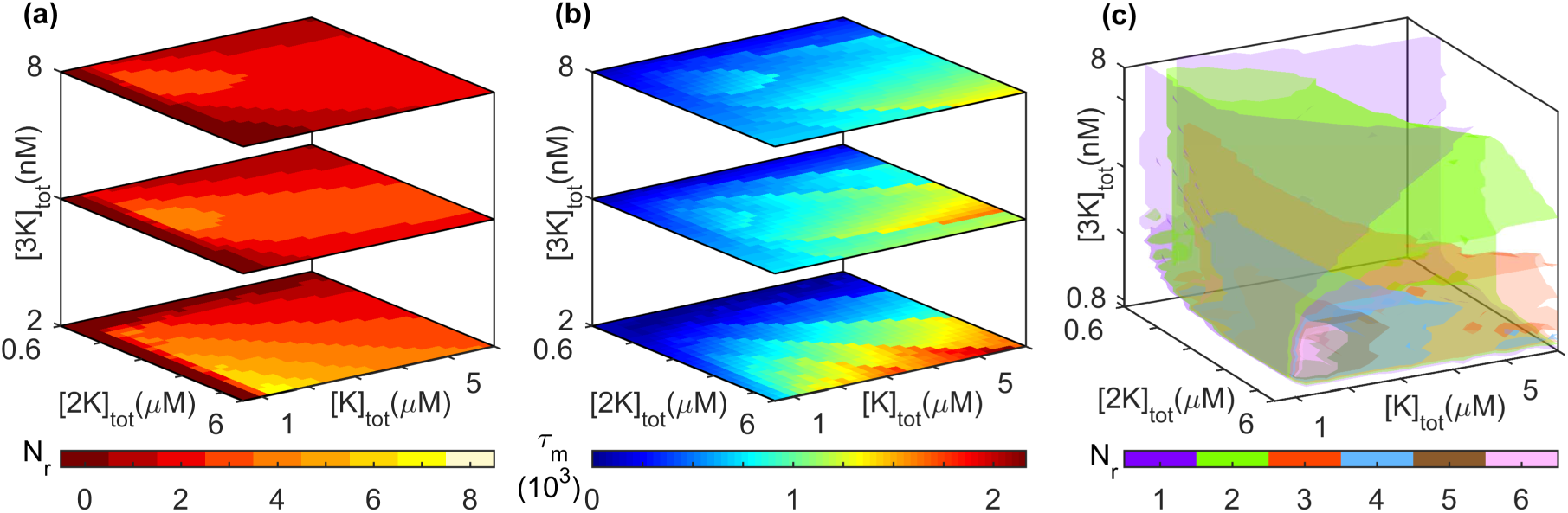
Dependence of reverberatory activity on the total kinase concentrations, viz., MAPK ([*K*]_*tot*_), MAP2K ([2*K*]_*tot*_) and MAP3K ([3*K*]_*tot*_). (a) The number of post-stimulus spikes *N_r_*, (b) the total memory time *τ_m_* (in minutes), and (c) isosurfaces for *N_r_* observed on withdrawing an applied stimulus of amplitude S = 2.0 × 10^−6^*µM*, are shown as functions of the three total kinase concentrations. The total concentrations of the phosphatases are held fixed for (a-c) and are provided in Table S4.

**FIG. S5:**
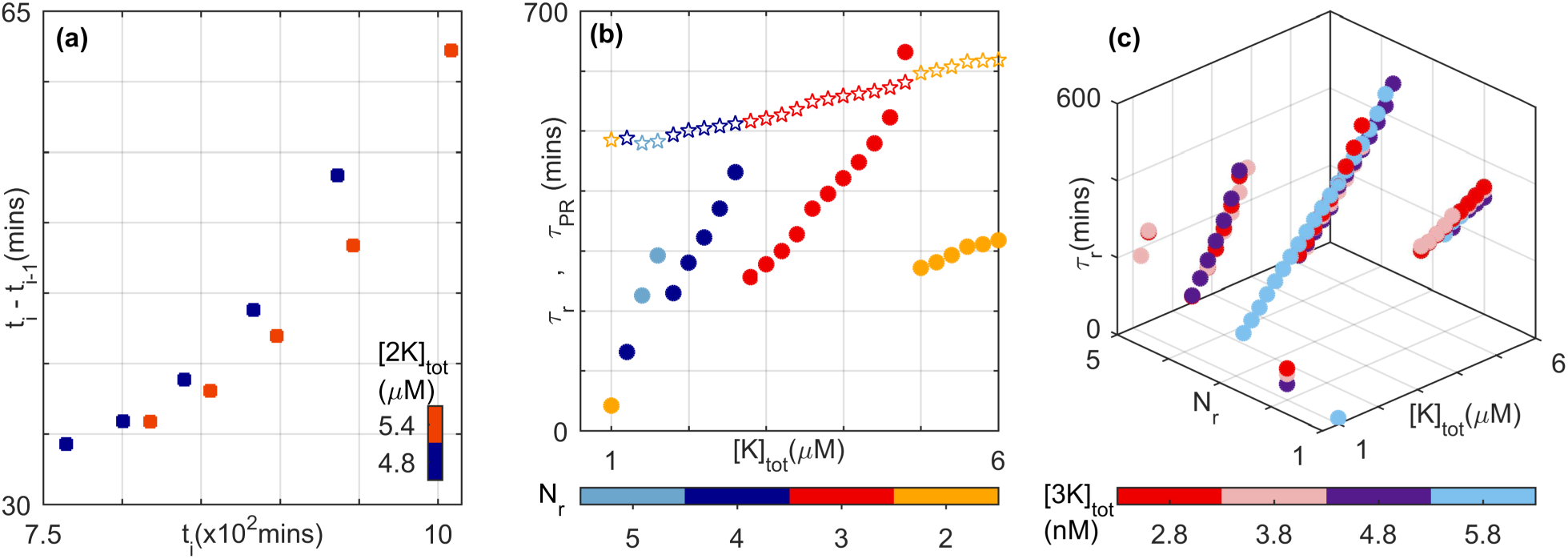
Characterization of the reverberatory dynamics observed after withdrawing a stimulus having amplitude *S* = 2.0 × 10^−6^ *µM*. (a) The interval between successive spikes *i* − 1 and *i* increases with time (*t_i_* being the time of occurrence of the ith spike) for two distinct total concentrations of MAP2K. The total concentrations of MAPK and MAP3K are [*K*]_*tot*_ = 1.2*µM* and [3*K*]_*tot*_ = 2.8nM, respectively. (b) The primary recovery time *τ_PR_* (stars) and the total duration of reverberatory activity *τ_r_*(filled circles) are shown for different values of *N_r_* (indicated by the color bar). While *τ_PR_* increases monotonically with increasing total MAPK concentration, *τ_r_*shows a more complex dependence ([2*K*]_*tot*_ = 3*µM* and [3*K*]_*tot*_ = 4nM). (c) The dependence of *τ_r_*on [*K*]_*tot*_ for different values of *N_r_* has a similar nature for different choices of [3*K*]_*tot*_ (indicated by the color bar, [2*K*]_*tot*_ = 3*µM*). Note that for panel (c), we consider only situations where the system attains a steady state on maintaining the stimulation. For the total concentrations of the phosphatases see Table S4.

**FIG. S6:**
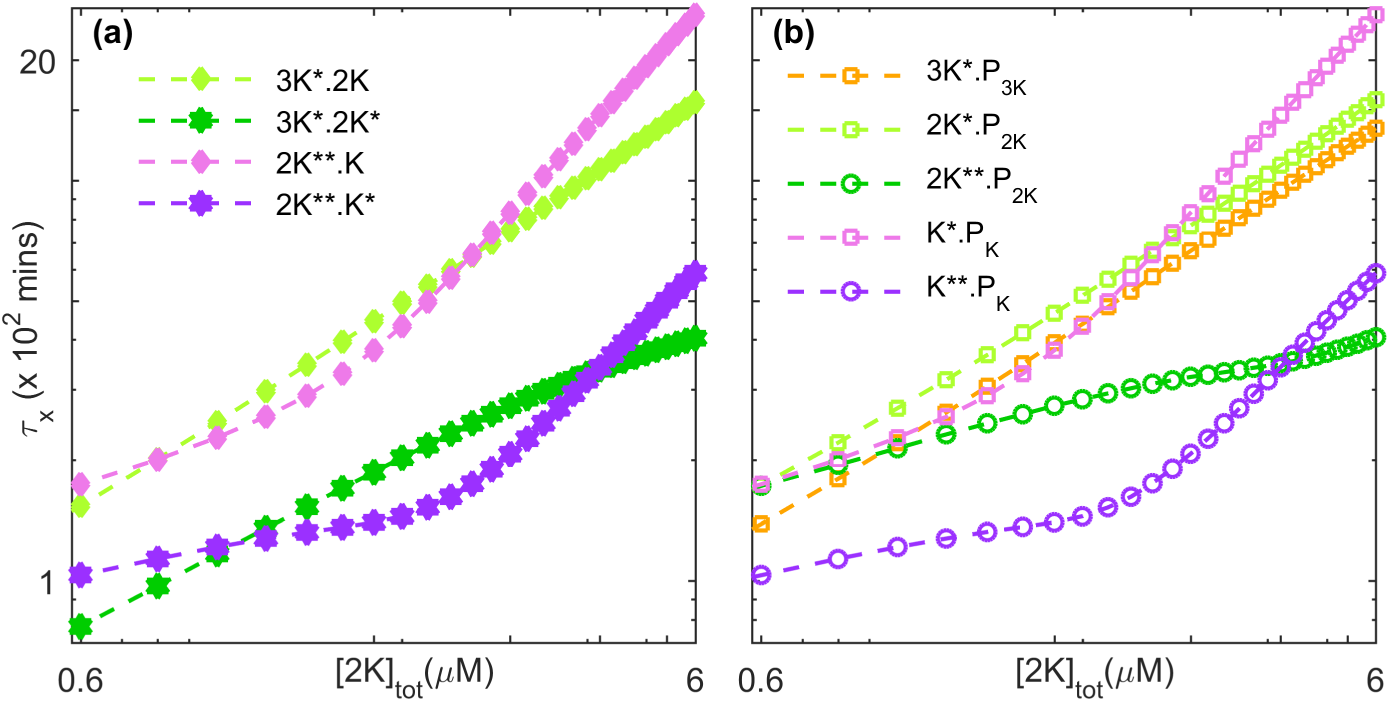
Protein complexes in the MAPK cascade exhibit relaxation behavior occurring over a broad range of time-scales. Decay of activity is shown after withdrawing an applied stimulus of amplitude *S* = 1.2×10^−6^ *µM*. The relaxation times *τ*_x_ of the different molecular species, viz., (a) the protein complexes between non-phosphorylated and singly phosphorylated (non-active) kinase proteins and the doubly phosphorylated (active) kinase protein of the preceding layer, and (b) the protein complexes between the phosphorylated (singly-or doubly-) kinase proteins and the phosphatase that carries out dephosphorylation in the corresponding layer of the MAPK cascade, vary with the total concentration of MAP2K. The nature of this dependence is distinct for lower and higher values of [2*K*]_*tot*_. For both panels, [*K*]_*tot*_ = 0.8*µM* and [3*K*]_*tot*_ = 0.0020*µM*. The total concentrations of the phosphatases are provided in Table S4.

**FIG. S7:**
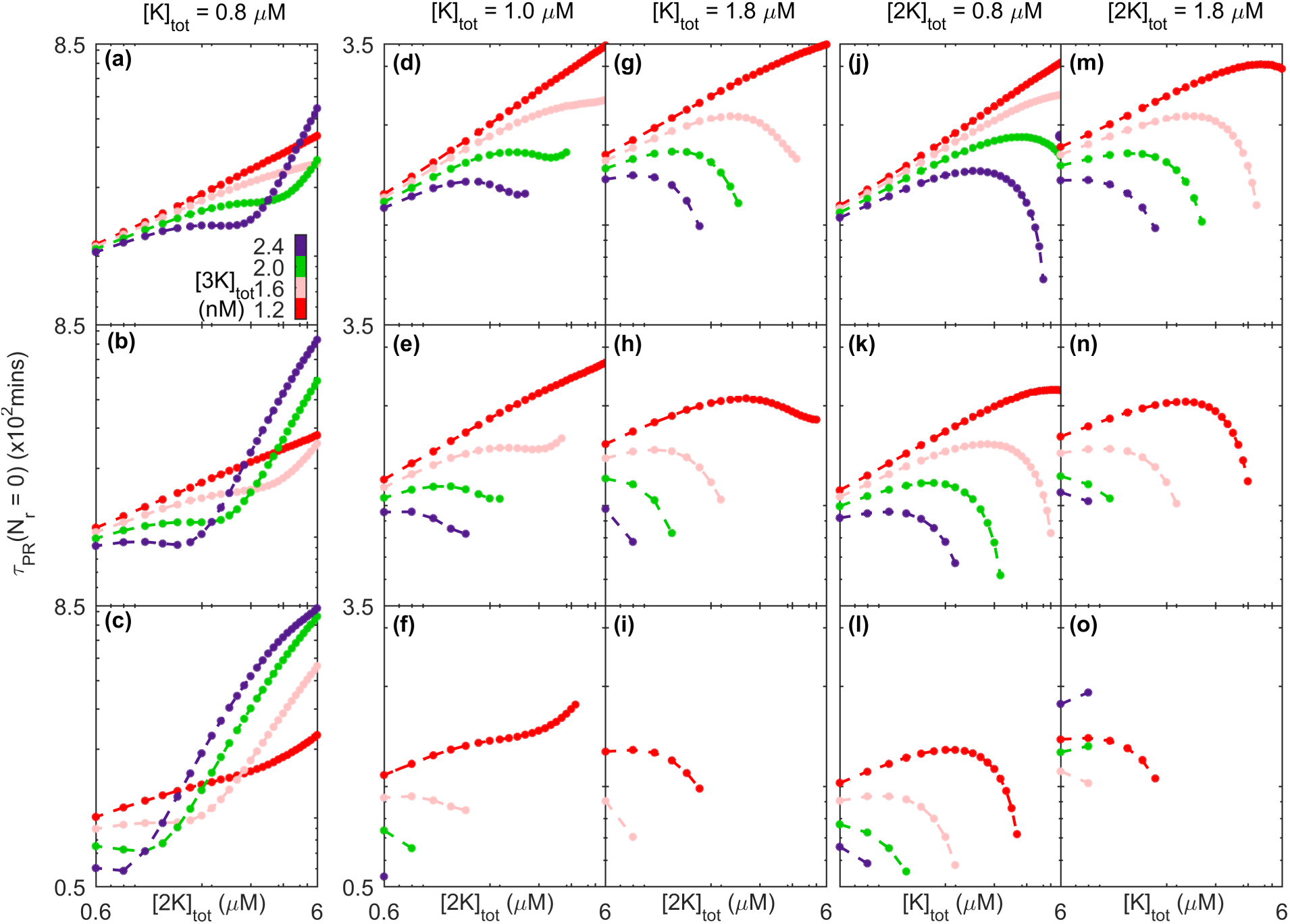
Dependence of the primary recovery time *τ_PR_* on (a-i) the total concentration of MAP2K ([2*K*]_*tot*_) and on (j-o) the total concentration of MAPK ([*K*]_*tot*_) for different values of the total concentration of MAP3K ([3*K*]_*tot*_), obtained upon removing stimuli having different amplitudes S. Panels (a,d,g,j,m) are for *S* = 0.8 × 10^−6^*µM*, panels (b,e,h,k,n) are for *S* = 1.2 × 10^−6^ *µM*, and panels (c,f,i,l,o) are for *S* = 2.0 × 10^−6^ *µM*. We have only considered situations where the system reaches a steady state upon application of a time-invariant stimulus, and that do not show any reverberatory activity (*N_r_* = 0) during relaxation to the resting state. The curves in panels (a-i) are obtained for different values of [*K*]_*tot*_, namely, (a-c) [*K*]_*tot*_ = 0.8*µM*, (d-f) [*K*]_*tot*_ = 1.0*µM*, and (g-i) [*K*]_*tot*_ = 1.8*µM*. The curves in panels (j-o) are obtained for different values of [2*K*]_*tot*_, namely, (j-l) [2*K*]_*tot*_ = 0.8*µM*, and (m-o) [2*K*]_*tot*_ = 1.8*µM*. The total concentrations of the phosphatases for all panels are given in Table S4.

**FIG. S8:**
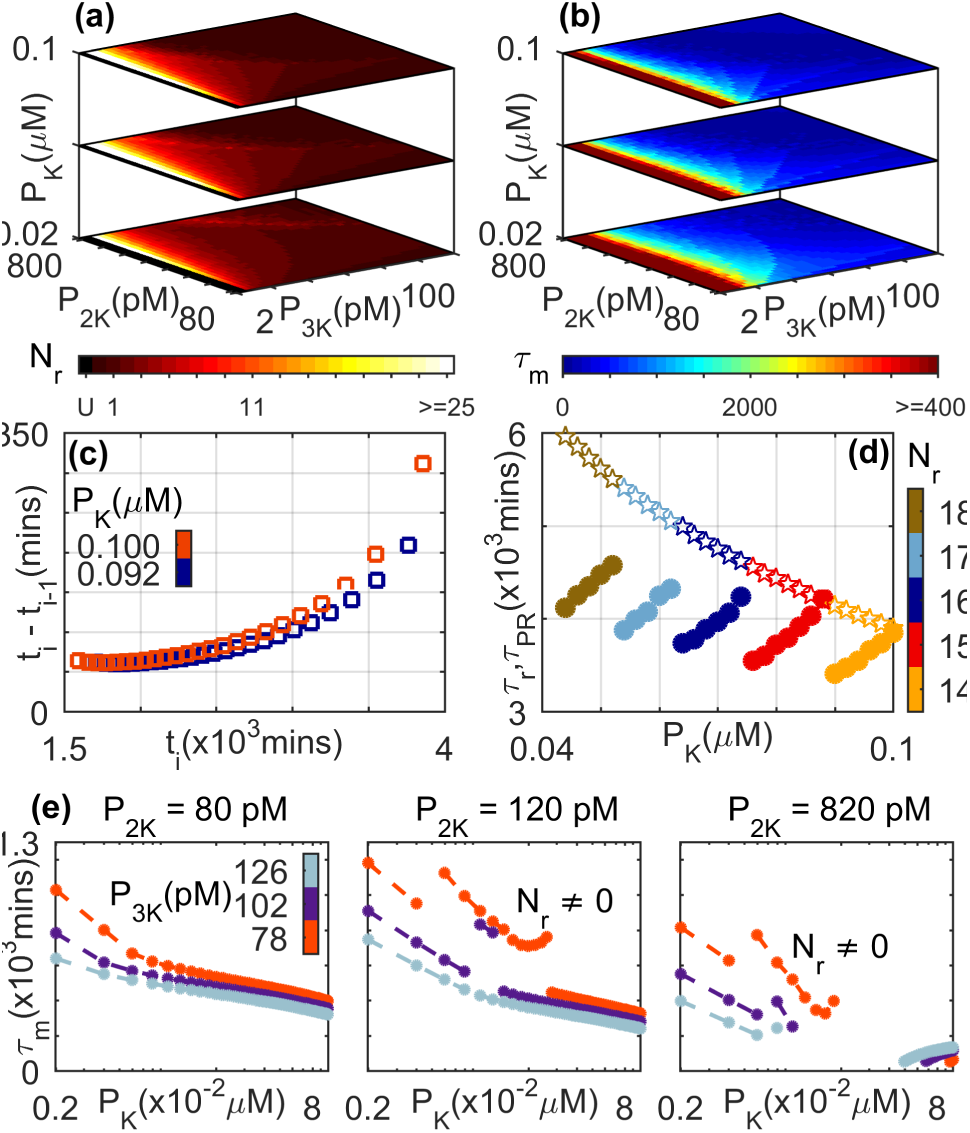
Dependence of reverberatory activity on the total concentrations of the phosphatases MAPK P’ase ([*P_K_*]), MAP2K P’ase, ([*P_2K_*]) and MAP3K P’ase ([*P_3K_*]). (a) The number of spikes *N_r_* and (b) the total memory time *τ_m_* (in minutes) observed on withdrawing an applied stimulus of amplitude *S* = 2.0 × 10^−6^*µM*. Situations where the primary recovery time is longer than a maximum or cut-off value (see Methods), such that the reverberatory nature of the dynamics cannot be properly measured, are indicated by the color corresponding to “U”. (c) The interval between successive spikes *i* − 1 and *i* increases with time (*t_i_* being the time of occurrence of the ith spike). As the MAPK P’ase concentration is increased, the durations of these intervals are seen to increase. The total concentrations of the other two phosphatases are maintained at [*P_2K_*] = 680*pM* and [*P_3K_*] = 10*pM*. (d) The variation of primary recovery time *τ_PR_* (stars) and the total duration of reverberatory activity *τ_r_*(filled circles) as a function of total MAPK P’ase concentration are shown for different values of *N_r_* (indicated by the color bar). While *τ_PR_* decreases monotonically with increasing [*P_K_*], *τ_r_*shows a more complex dependence ([*P_2K_*] = 200*pM* and [*P_3K_*] = 6*pM*). (e) Dependence of the total memory time *τ_m_* on total MAPK P’ase concentration ([*P_K_*] shown in log scale) for different total concentrations of MAP2K P’ase (values indicated above each of the three panels) and MAP3K P’ase (indicated using different colors as shown in the color bar). Note that we consider only situations where the system attains a steady state on maintaining stimulation. For details of the total concentrations of the kinases, see Table S3.

**FIG. S9:**
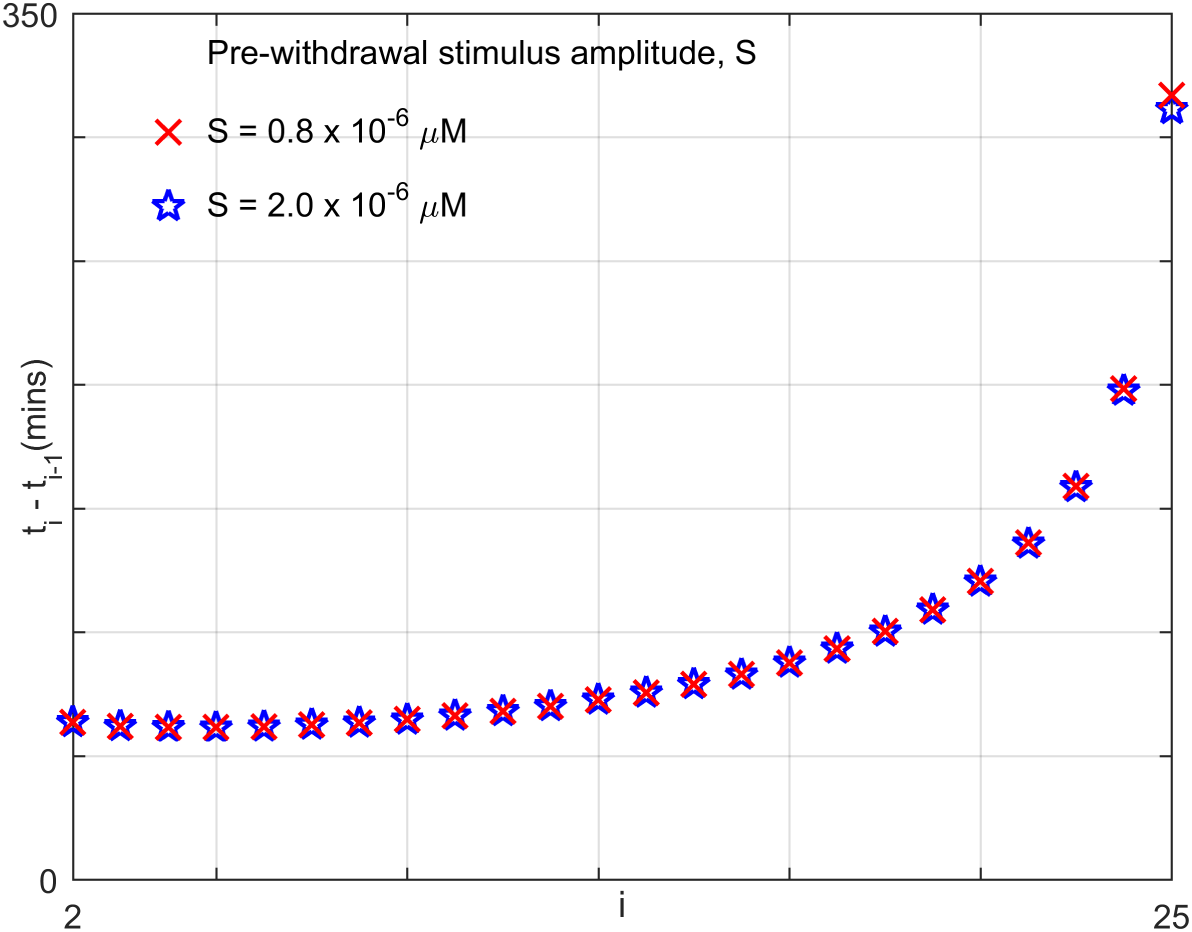
The time interval between successive spikes *i* − 1 and *i* obtained after removing a stimulus, increases with the number of spike events (*i* being the event number of the ith spike). The trend appears to be independent of the stimulus amplitude S. The total concentrations of the phosphatases are *P_K_*= 0.1*µM*, *P_2K_*= 680*pM* and *P_3K_* = 10*pM*, respectively. The total concentrations of the kinases are provided in Table S3.

## References

[1] U. S. Bhalla and R. Iyengar, Science 283, 381 (1999).

[2] J. E. Purvis and G. Lahav, Cell 152, 945 (2013).

[3] B. N. Kholodenko, Nat. Rev. Mol. Cell Bio. 7, 165 (2006).

[4] G. Weng, U. S. Bhalla, and Iyengar R., Science 284, 92 (1999).

[5] C. Widmann, S. Gibson, M. B. Jarpe, and G. L. Johnson, Physiol. Rev. 79, 143 (1999).

[6] R. Seger and E. G. Krebs, FASEB J. 9, 726 (1995).

[7] A. J. Waskiewicz and J. A. Cooper, Curr. Opin. Cell Biol. 7, 798 (1995).

[8] B. Su and M. Karin, Curr. Opin. Immunol. 8, 402 (1996).

[9] B. Alberts, A. Johnson, J. Lewis, D. Morgan, M. Raff, K. Roberts, and P. Walter, Molecular Biology of the Cell, 6th Ed. (Garland Science, New York, 2014).

[10] C. Y. Huang and J. E. Ferrell, Proc. Natl. Acad. Sci. USA 93, 10078 (1996).

[11] H. Shankaran, D. L. Ippolito, W. B. Chrisler, H. Resat, N. Bollinger, L. K. Opresko, and H. S. Wiley, Mol. Sys. Biol. 5, 332 (2009).

[12] L. Qiao, R. B. Nachbar, I. G. Kevrekidis, and S. Y. Shvartsman, PLOS Comput. Biol. 3, e184 (2007).

[13] T. Jesan, U. Sarma, S. Halder, B. Saha, and S. Sinha, PLOS One 8, e64409 (2013).

[14] A. C. Ventura, J. A. Sepulchre, and S. D. Merajver, PLOS Comput. Biol. 4, e1000041 (2008).

[15] N. I. Markevich, J. B. Hoek, and B. N. Kholodenko, J. Cell Biol. 164, 353 (2004).

[16] S. Legewie, B. Schoeberl, N. Blüthgen, and H. Herzel, Biophys. J. 93, 2279 (2007).

[17] F. Ortega, J. L. Garcés, F. Mas, B. N. Kholodenko, and M. Cascante, FEBS J. 273, 3915 (2006).

[18] J. E. Ferrell, Curr. Opin. Cell Biol. 14, 140 (2002).

[19] M. C. Inniss and P. A. Silver, Curr. Biol. 23, R812 (2013).

[20] W. Xiong and J. E. Ferrell, Nature (London) 426, 460 (2003).

[21] C. M. Ajo-Franklin, D. A. Drubin, J. A. Eskin, E. P. Gee, D. Landgraf, I. Phillips, and P. A. Silver, Gene. Dev. 21, 2271 (2007).

[22] D. R. Burrill and P. A. Silver, Cell 140, 13 (2010).

[23] J. Gunawardena, Proc. Natl. Acad. Sci. USA 102, 14617 (2005).

[24] O. Hadac, I. Schreiber, and M. Pribyl, J. Chem. Phys. 138, 065102 (2013).

[25] G. T. Philips, X. Ye, A. M. Kopec, and T. J. Carew, J. Neurosci. 33, 7565 (2013).

[26] G. Y. Wu, K. Deisseroth, and R. W. Tsien, Nat. Neurosci. 4, 151 (2001).

[27] J. D. Sweatt, J. Neurochem. 76, 1 (2001).

[28] C. Huang, K. Jacobson, and M. D. Schaller, J. Cell Sci. 117, 4619 (2004).

[29] C. Dong, R. J. Davis, and R. A. Flavell, Ann. Rev. Im-munol. 20, 55 (2002).

[30] G. Altan-Bonnet and R. N. Germain, PLOS Biol. 3, e356 (2005).

[31] T. Millat, E. Bullinger, J. Rohwer, and O. Wolkenhauer, Math. Biosci. 207, 40 (2007).

[32] S. Filippi, C. P. Barnes, P. D. Kirk, T. Kudo, K. Kunida, S. S. McMahon, T. Tsuchiya, T. Wada, S. Kuroda, and M. P. Stumpf, Cell Rep. 15, 2524 (2016).

[33] M. Jeschke, S. Baumg¨artner, and S. Legewie, PLOS Comput. Biol. 9, e1003357 (2013).

[34] S. Mukherjee, S. C. Seok, V. J. Vieland, and J. Das, Proc. Natl. Acad. Sci. USA 110, 18531 (2013).

[35] C. J. Marshall, Cell 80, 179 (1995).

[36] A. Plotnikov, E. Zehorai, S. Procaccia, and R. Seger, BBA-Mol. Cell Res. 1813, 1619 (2011).

[37] Q. Lu, M. Paredes, J. Zhang, and K. S. Kosik, Mol. Cell. Biol. 18, 3257 (1998).

[38] X. Ye, J. L. Shobe, S. K. Sharma, A. Marina, and T. J. Carew, Proc. Natl. Acad. Sci. USA 105, 20511 (2008).

[39] C. M. Atkins, J. C. Selcher, J. J. Petraitis, J. M. Trza-skos, and J. D. Sweatt, Nat. Neurosci. 1, 602 (1998).

[40] M. Skoge, H. Yue, M. Erickstad, A. Bae, H. Levine, A. Groisman, W. F. Loomis, and W. J. Rappel, Proc. Natl. Acad. Sci. USA 111, 14448 (2014)

[41] H. V. Prentice-Mott, Y. Meroz, A. Carlson, M. A. Levine, M. W. Davidson, D. Irimia, G. T. Charras, L. Mahade-van, and J. V. Shah, Proc. Natl. Acad. Sci. USA 113, 1267 (2016).

[42] X. Guo, L. Wang, B. Chen, Q. Li, J. Wang, M. Zhao, W. Wu, P. Zhu, X. Huang, and Huang Q., Am. J. Physiol.-Heart C. 297, H238 (2009).

[43] M. Koss, G. R. Pfeiffer, Y. Wang, S. T. Thomas, M. Yerukhimovich, W. A. Gaarde, C. M. Doerschuk, and Q. Wang, J. Immunol. 176, 1218 (2006).

